# High cryptic diversity of invertebrates in varying nearshore habitats of the Beaufort Sea

**DOI:** 10.64898/2026.09.08.750166

**Authors:** Sabrina Heiser, Lorine Salel, Susan Schonberg, Deana L. Erdner, Kenneth H. Dunton

**Author notes:** Corresponding author: Sabrina Heiser, Marine Science Institute, University of Texas at Austin, 750 Channel View Dr., Port Aransas, Texas, 78373, United States.

## Abstract

The Alaskan Beaufort Sea lagoon system supports a diverse invertebrate community shaped by a complex glacial and biogeographic history, yet it remains severely understudied relative to the rapid environmental changes it is experiencing. As Arctic amplification accelerates habitat loss, shifts dispersal pathways, and facilitates the introduction of boreal and invasive species, accurately documenting current biodiversity is increasingly urgent. In this study, we used DNA barcoding of the mitochondrial *cox*1 gene for various invertebrates, and the nuclear 28S rRNA gene for Porifera, to examine invertebrate genera across coastal sites in the Alaskan Beaufort Sea, including the unique Boulder Patch kelp community in Stefansson Sound. We sequenced specimens belonging to five phyla: Annelida, Arthropoda, Mollusca, Porifera, and Priapulida. Specimens were identified by morphology, and their *cox*1 or 28S rRNA gene sequences were matched against the GenBank and Barcode of Life Data Systems (BOLD) databases. Molecular identification revealed substantially higher species diversity than morphology alone in three of five phyla: Annelida (12 versus six species), Arthropoda (14 versus 11), and Mollusca (five versus four). In Porifera and Priapulida the number of species were the same for morphological and molecular identification; however, we still found different species identifications between the two methods. Other notable findings include the discovery of multiple cryptic species within *Terebellides* sp., *Pontoporeia femorata*, *Onisimus litoralis*, and *Micronephthys minuta*, as well as the first published sequences for *Acanthostepheia incarinata*, *Haliclona gracilis*, *Onisimus affinis*, and *Saduria sibirica*. Our results highlight the extent to which morphology-based surveys underestimate invertebrate diversity in Arctic ecosystems and underscore the need for comprehensive surveys and molecular reference databases to support future molecular-based biodiversity monitoring in this rapidly changing region.

## Introduction

Rapid environmental change is transforming Arctic coastal ecosystems faster than they can be documented, creating an urgent need to establish biodiversity baselines before these communities are fundamentally altered (Deb & Bailey, 2023). The Arctic is experiencing climate change much faster than the global average, a phenomenon known as Arctic amplification, with the poles warming approximately four times faster than the rest of the world (Rantanen et al., 2022; Serreze & Barry, 2011). Rising seawater temperatures and declining sea ice coverage are modifying habitat suitability and dispersal pathways (Hardy et al., 2011) including increasing species transport from the Pacific to the Arctic (Watanabe et al., 2014). The introduction of new species from the expansion of boreal ranges into Arctic ecosystems and invasive species via newly opened shipping routes as sea ice retreats have profound consequences for Arctic biodiversity (Descamps et al., 2026; Ware et al., 2016). Together, these changes introduce new competitors that place increasing pressure on native Arctic species. Furthermore, as climate change continues to reshape the Arctic environment, invasive species may hold a competitive advantage over native fauna that may lack the physiological flexibility to adapt and compete in the changing climate, therefore raising concerns about long-term biodiversity loss (Callaghan et al., 2004; Descamps et al., 2026; Lemieux et al., 2025). Thoroughly documenting current biodiversity is therefore essential for detecting these changes and understanding what is at risk (Bluhm et al., 2011).

Establishing an accurate species inventory for the Arctic is complicated by the region’s complex glacial and biogeographic history. The geological events of the Pleistocene and the subsequent recolonizations have produced communities that are strongly influenced by both Pacific and Atlantic taxa and exhibit high cryptic diversity across many taxonomic groups (Bringloe, Verbruggen, & Saunders, 2020; Dunton, 1992). The first opening of the Bering Strait during the Pliocene, approximately 4.8–5.5 Ma, had a significant impact on Arctic marine fauna by establishing a connection between the Arctic and Pacific basins. Initially, ocean circulation through the strait facilitated the movement of Atlantic taxa through the Arctic and into the Pacific. Later, the progressive restriction of the Central American Seaway associated with the formation of the Isthmus of Panama altered ocean currents around North America. By approximately 3.6 Ma, flow through the Bering Strait had shifted toward the modern Pacific-to-Arctic direction, facilitating the dispersal of Pacific taxa into Arctic and Atlantic waters (Bringloe & Saunders, 2019; Haug & Tiedemann, 1998; Marincovich, 2000). During the Pleistocene, alternating glacial cycles subsequently caused periods of isolation, local extinction, and recolonization of Arctic marine fauna. These repeated episodes of glaciation contributed to the relatively young and complex evolutionary history of many Arctic taxa (Dunton, 1992; Väinölä, 2003). Additionally, extreme environments, such as found in the polar regions, have been postulated to support morphological stasis during speciation (Bickford et al., 2007). As a result, many of these sister taxa, specifically in invertebrates, are morphologically indistinguishable and likely harbor far more cryptic diversity than morphological differences alone can reveal (Laakkonen, Hardman, Strelkov, & Väinölä, 2021).

In the Alaskan Beaufort Sea lagoons, this already complicated evolutionary history is compounded by highly variable environmental conditions. The lagoons are shallow and vary in resource availability, yet they support a diverse and resilient benthic fauna that endures extremes, such as periods of hypoxia, hyper- and hyposalinity, as well as intense periods of seasonal darkness and sea ice cover (Dunton et al., 2019). The high seasonal variability and restricted exchange with oceanic waters in the lagoons create systems that are ecologically dynamic but also highly sensitive to environmental change, making them a priority for long-term biodiversity monitoring. This study sampled lagoon sites from the far west to the far east of the Alaskan Beaufort Sea coastline, with differing environmental conditions and connections to the Beaufort Sea (Figure 1). A highlight of our sampling is the Stefansson Sound Boulder Patch, an area of hard stratum supporting the richest and most biologically diverse benthic community discovered to date in the Alaskan Beaufort Sea (Dunton, Reimnitz, & Schonberg, 1982; Dunton & Schonberg, 2000). The diverse site locations make Arctic marine invertebrates fascinating for studying speciation but challenging for traditional morphology-based taxonomy, particularly in a region as biogeographically complex as the Alaskan Beaufort Sea. Despite decades of morphological surveys, the invertebrate diversity of both types of habitats (hard substratum and soft bottom) has rarely been characterized using a molecular approach to aid species delimitation.

**Figure 1.**
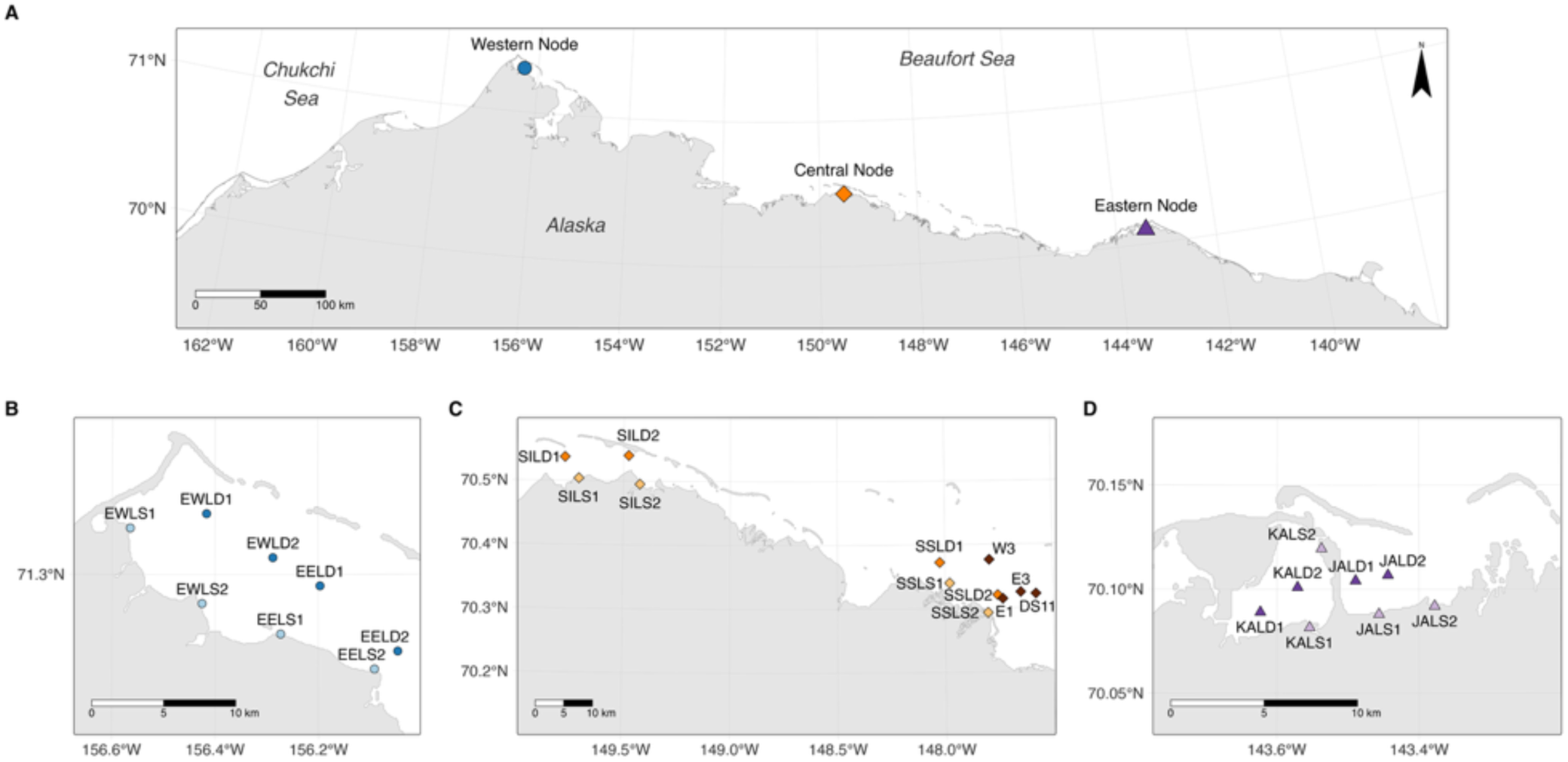
(A) Location of sampling sites along the northern slope of Alaska including the (B) western node (blu western (EWL) and eastern (EEL) area of Elson lagoon, (C) central node (orange squares) with Simpson lagoon Sound (SSL) and Boulder Patch sites (brown squares; W3, E1, E3, DS11), and (D) eastern node (purple triangles (KAL) and Jago (JAL) lagoons. Lighter shades indicate shallow and darker shades indicate deeper sites which w trawl or Ponar grab, except for the Boulder Patch which was sampled using SCUBA.

Molecular tools, particularly DNA barcoding, help to accurately characterize biodiversity, specifically when species are too morphologically similar to distinguish reliably or when juveniles lack identifying morphological features (Hardy et al., 2011). DNA barcoding addresses these limitations by providing genetic fingerprints for species, allowing not only the identification of unknown specimens but also the detection of some intraspecific genetic divergences (Hebert, Ratnasingham, & deWaard, 2003; Walczyńska, Mańko, & Weydmann, 2018). Barcoding facilitates the recognition of cryptic species with implications for how we identify threatened or endangered species and how we assess biodiversity loss, as cryptic species may differ in ecological function (Fišer, Altermatt, Zakšek, Knapič, & Fišer, 2015). Furthermore, barcodes submitted to a publicly accessible database such as GenBank (NCBI) and the Barcode of Life Data Systems (BOLD) build reference databases that facilitate data sharing and support the use of powerful emerging tools such as eDNA metabarcoding (Cristescu, 2014; Hardy et al., 2011) which, unlike single-specimen barcoding, expands this approach to the entire community, enabling the identification of multiple species from a single sample (Ji et al., 2013).

Here, we use DNA barcoding to characterize the benthic invertebrate communities across several types of habitats of the Alaskan Beaufort Sea and identify cryptic diversity within species identified by morphology (morphospecies). In doing so, we identify higher than expected species diversity in three out of five phyla and contribute new reference sequences for previously un-sequenced taxa to public databases.

## Methods

### Sample collection

Invertebrates were collected from three nodes along the Beaufort Sea lagoonal system and the Boulder Patch in Stefansson Sound (Figure 1). Sites were generally identified using a three-letter code followed by a fourth letter indicating depth (S=shallow, D=deep), and a number. Some samples lack a numerical designation because they were collected near, but not directly at the predefined coordinates. The four Boulder Patch sites within Stefansson Sound were labeled separately (Figure 1).

Samples were collected using SCUBA in the Boulder Patch and with Ponar grabs and bottom trawls at lagoon sites (see Supplemental Table 1 for details). Ponar grab samples were collected at shallow stations ranging from 0.64 to 1.81 m depth and at deep stations ranging from 2.40 to 3.70 m depth. Specimens were preserved in 100 % (200-proof) ethanol and transported to the University of Texas at Austin Marine Science Institute (UTMSI), where they were identified morphologically by Susan Schonberg and archived in the Dr. Kenneth H. Dunton invertebrate collection.

Maps of sampling locations were produced in R version 4.4.0 (R Core Team, 2024) using the packages *cowplot* (Wilke, 2025), *grid* (R Core Team, 2024), *png* (Urbanek, 2022), *ragg* (Pedersen & Shemanarev, 2025), *sf* (Pebesma, 2018; Pebesma & Bivand, 2023), and *tmap* (Tennekes, 2018). The Global Self-consistent Hierarchical High-resolution Geography (GSHHG) database version 2.3.7 served as the basemap (Wessel & Smith, 1996).

### DNA extraction

For most specimens, genomic DNA was extracted from small tissue fragments (≤ 8 mm³) using an adapted cetyltrimethylammonium bromide (CTAB) based protocol (Panova et al., 2016, Supplemental Table 1 for details on tissue type). Dissected tissues were placed in 500 µL CTAB lysis buffer (10450002-1, BioWorld, Dublin, OH, USA) with 1 µL β-mercaptoethanol (35602BID, Thermo Scientific, Rockford, IL, USA) and 20 µL Proteinase K (EO0492, Thermo Fisher Scientific Inc, Nashville, Tennessee, USA) and incubated at 60°C on a shaking platform for one hour. Samples were then mechanically homogenized using one bead and a vortex for five minutes, followed by an additional 30-minute incubation at 60°C to allow foam to dissipate.

RNA was removed by adding 5 µL RNase A (EN0531, Thermo Fisher Scientific), followed by brief vortexing, and then incubating for one hour at 60°C. Phase separation was performed by adding 500 µL of 24:1 chloroform:isoamyl alcohol mixture prepared from chloroform (MP Biomedicals, Solon, OH, USA) and isoamyl alcohol (Fisher Scientific, Waltham, MA, USA) and then gently inverting for two minutes, followed by centrifugation at 14,000×g at 4°C for 10 minutes. Approximately 390 µL of the aqueous phase was transferred to a clean 1.5 mL tube, and DNA was precipitated overnight at room temperature by adding 260 µL isopropanol (A417-1, Fisher Chemical, Fair Lawn, NJ, USA). DNA was pelleted at 14,000×g at 4°C for 30 minutes the next day. The supernatant was removed, and the pellet was washed with 600 µL ice-cold 70% ethanol (Fisher BioReagents, Waltham, MA, USA), air-dried, and resuspended in 20 µL sterile nuclease-free water (D4302-5-10, Zymo Research, Irvine, CA, USA).

For some groups of specimens that were consistently unsuccessful with the CTAB protocol, DNA was extracted using the DNeasy Blood & Tissue Kit (Qiagen) following the manufacturer’s protocol. Some DNA extracts were re-extracted using CTAB to reduce contaminants. Tissue selection and cleaning varied by phylum (Supplemental Table 1).

The DNA was quantified using the AccuBlue High Sensitivity dsDNA Quantification Kit (99940, Biotium, Fremont, California, USA) and a CLARIOstar plate reader (BMG Labtech, Cary, North Carolina, USA). We then stored the DNA at 4°C until PCR amplification for short term and at -20°C for longer term (>1 week).

### PCR amplification

Polymerase Chain Reaction (PCR) amplifications consisted of two 25 µL reactions: an initial reaction using 150 ng of DNA, followed by a second 25 µL reaction if amplification was successful. For specimens that did not amplify, reactions were repeated using 25 ng of DNA (see Supplemental Table 1 for more details on DNA concentrations for each sample).

We performed PCR amplifications for the mitochondrial *cox*1 gene and the nuclear 28S rRNA gene using DreamTaq DNA Polymerase (EP0702, Thermo Fisher Scientific). Each 25 µL reaction contained 2.5 µL DreamTaq buffer, 2.5 µL of 2 mM of each dNTP, 1.5 µL of each forward and reverse primer (at 10 µM each), 0.2 µL DreamTaq DNA Polymerase, and template DNA, following the manufacturer’s protocol.

The mitochondrial *cox*1 gene was amplified using the universal invertebrate barcoding primers jgLCO1490 (5′-TITCIACIAAYCAYAARGAYATTGG-3′) and jgHCO2198 (5′-TAIACYTCIGGRTGICCRAARAAYCA-3′) generating an amplicon ∼700 bp long (Folmer, Black, Hoeh, Lutz, & Vrijenhoek, 1994; Geller, Meyer, Parker, & Hawk, 2013). The temperature profile consisted of an initial denaturation at 95°C for 3 minutes, followed by 35 cycles of 95°C for 30 seconds, 48°C for 30 seconds, and 72°C for 1 minute, with a final extension at 72°C for 10 minutes which was done using a Mastercycler nexus gradient thermal cycler (Eppendorf AG, Hamburg, Germany).

Specimens of Porifera amplified more reliably using the nuclear 28S rRNA gene with primers Por28S-15F (5’- GCGAGATCACCYGCTGAAT -3’) and Por28S-878R (5’-CACTCCTTGGTCCGTGTTTC -3’) generating an amplicon ∼800 bp long (Morrow et al., 2012). The temperature profile consisted of an initial denaturation at 94°C for 5 minutes, followed by 30 cycles of 94°C for 30 seconds, 53°C for 30 seconds, and 72°C for 1 minute, with a final extension at 72°C for 5 minutes.

We confirmed PCR amplification success using 0.7% agarose gel electrophoresis and performed a second round of PCR for samples that had faint or dark bands. PCR products were purified using the QIAquick PCR Purification Kit (28506, QIAGEN, Germantown, Maryland, USA), the MinElute PCR Purification Kit (28004, QIAGEN, Germantown, Maryland, USA), or the CleanPCR bead cleanup kit (CPCR-0050, CleanNA, Waddinxveen, Netherlands), following the manufacturers’ protocols.

### Sequencing and processing

The purified PCR products were sent to the Texas A&M University Corpus Christi (TAMUCC) Genomics Core Laboratory for Sanger sequencing. The resulting forward and reverse sequences were edited and aligned to generate a consensus using Geneious 2026.0.2 software. The consensus sequences were compared against reference sequences in GenBank and BOLD databases for taxonomic identification. Species were assigned at ≥98%, genetic identity (deWaard et al., 2019). If genetic identity was less than 98%, we inspected how the sequence clustered on the phylogenetic tree for taxonomy assignment. If any of our sequences constituted a novel addition to the public databases (i.e., no high genetic identity with any publicly available sequences and no entries under the morphospecies), we included the designation cf. (confer = compare with) in the species name we assigned to the sequence. Since we did not sequence the type specimens and cannot exclude the possibility of cryptic species, this is the most accurate name we can give the sequence based on the morphology of the specimen only. In some cases, our morphological identification for a specimen did not match the genetic species assignment based on sequences available in the databases. When our sequences matched a different species (≥98% genetic identity), we carefully consulted, where possible, the published literature associated with these sequences and usually reassigned the species names based on the genetic matches. When there were no matches with close genetic identity to any other sequence, including sequences submitted under the same species name, we included the designation aff. (affinis = related to) in the species name to signal a difference which we cannot resolve without access to type material.

We constructed phylogenetic trees for each phylum using sequences generated in this study and curated from the GenBank and BOLD databases (available sequences for species within the closest matching genus for each sequence). We used Geneious Prime with the Geneious Alignment (Global alignment with free end gaps) to align all sequences in the same taxonomic group and trim them to the same length. Maximum likelihood trees were constructed with IQ-TREE 3.0.1. We used ModelFinder to determine the best substitution model for each tree based on the Bayesian Information Criterion values (Kalyaanamoorthy, Minh, Wong, Von Haeseler, & Jermiin, 2017) and performed ultrafast bootstrap approximation with 1,000 replicates (Hoang, Chernomor, Von Haeseler, Minh, & Vinh, 2018). PearTree 1.3.1 (ARTIC Network) and Adobe Illustrator 2026 (Adobe Inc.) were used to prepare the final figures.

## Results

We extracted a total of 370 invertebrates belonging to approximately 50 different species based on morphological identification. From these extracts, we obtained 154 sequences corresponding to 30 species based on morphology and 41 species based on molecular analyses. All sequences (will) have been deposited in GenBank (Accession numbers: XXXX-XXXX) and BOLD (Dataset number: XXXX-XXX; see Supplemental Table 1 for details).

### Annelida

We successfully sequenced individuals identified as *Clymenura polaris* (n=1), *Embolocephalus nikolskyi* (n=1), *Laonome kroyeri* (n=2), *Micronephthys minuta* (n=8), *Nereimyra aphroditoides* (n=4), *Pista maculata* (n=1), *Pholoe* sp. (n=5), *Prionospio* sp. (n=2), and *Terebellides* sp. (n=13) based on morphology (Figure 2, Table 1). Overall intraspecific genetic identity ranged from 98.22-100% within Annelida.

**Figure 2.**
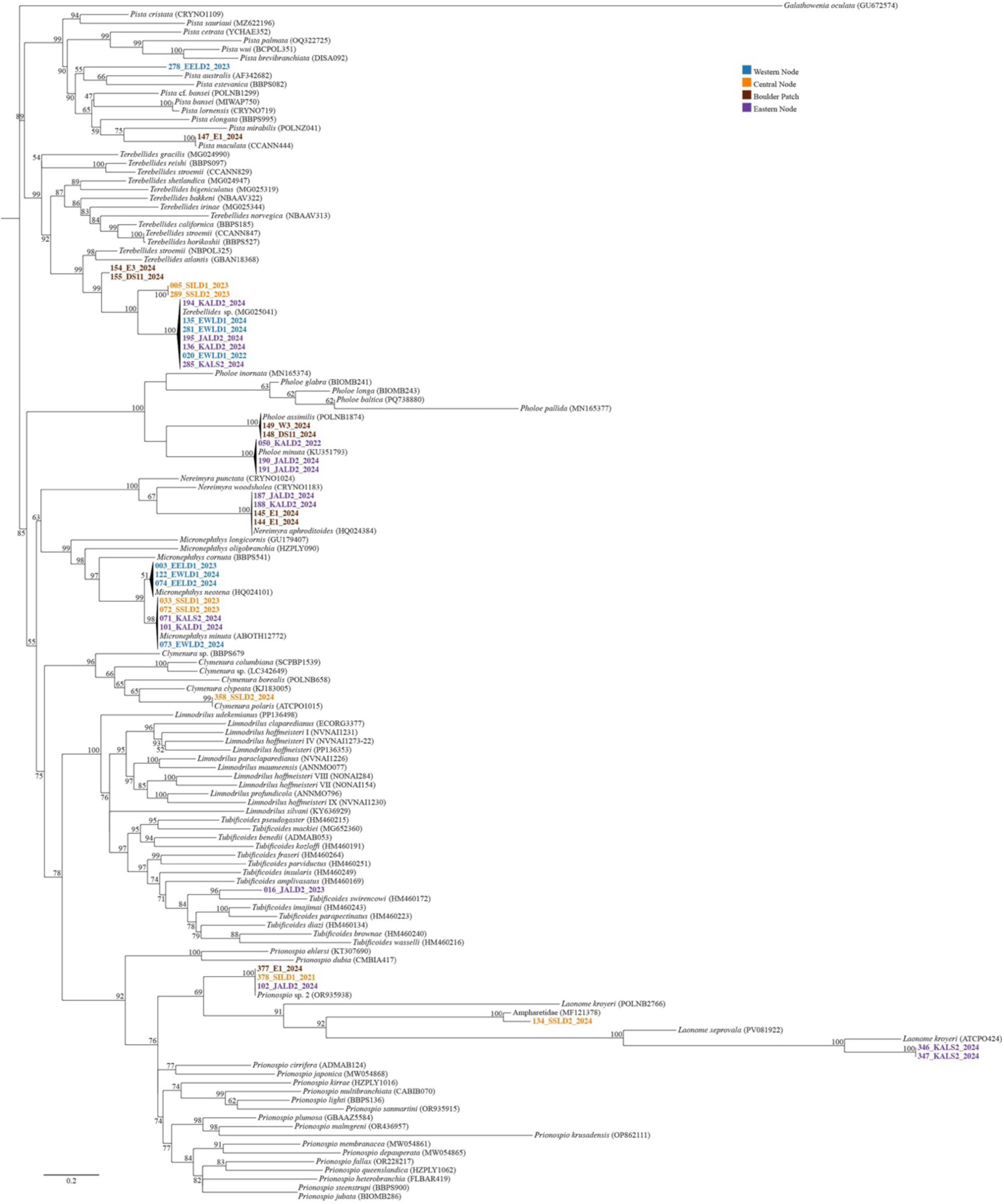
Maximum likelihood tree (*cox*1) for Annelida using the General Time Reversible substitution model with empirical base frequency, invariable sites, and the FreeRate Model with 5 categories (GTR+F+I+R5). Values indicate bootstrap support. *Galathowenia oculata* was used as the outgroup. Names in bold indicate sequences generated in this study with colors showing where they were collected from. All other sequences were sourced from the GenBank and BOLD databases (accession numbers provided).

**Table 1.**
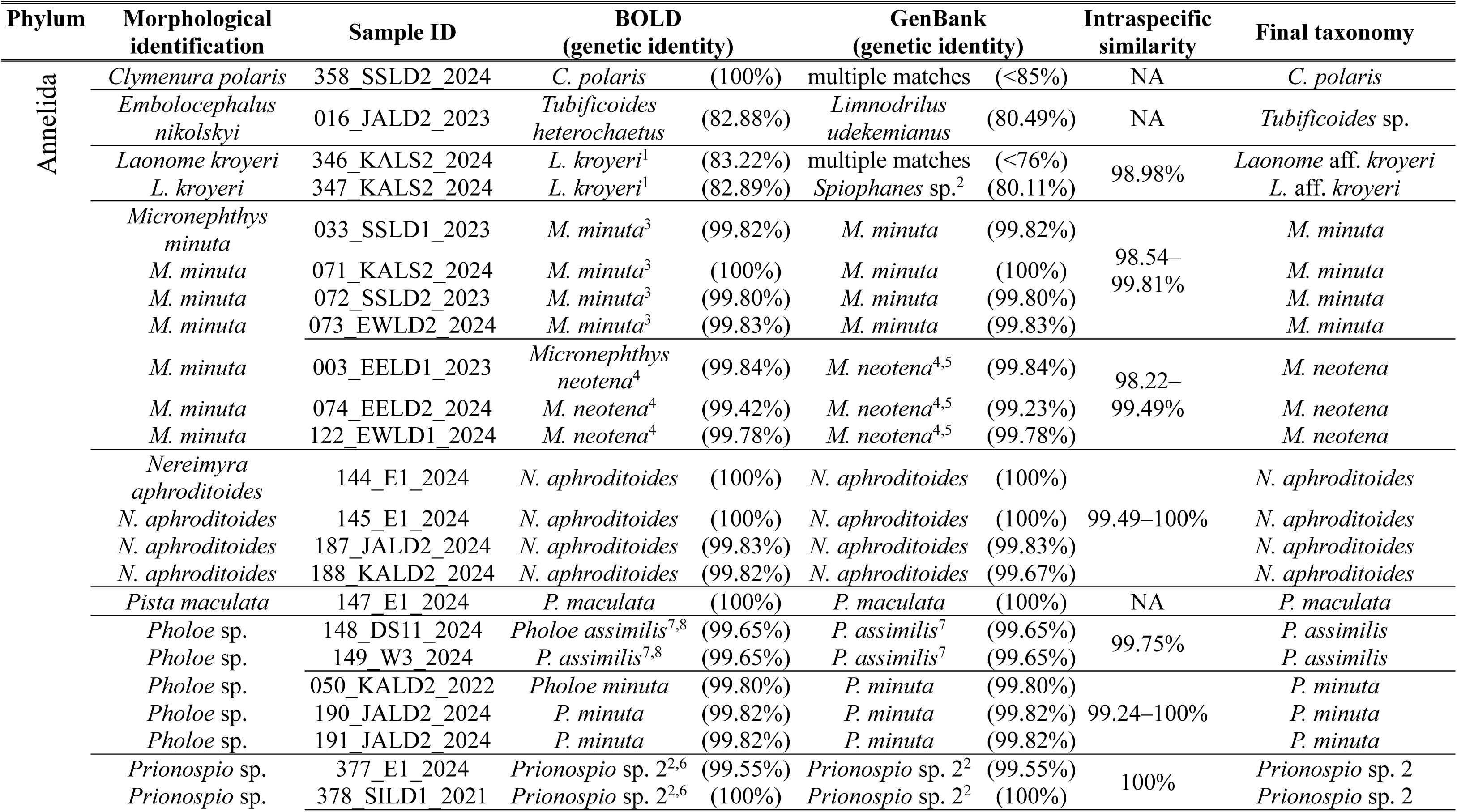
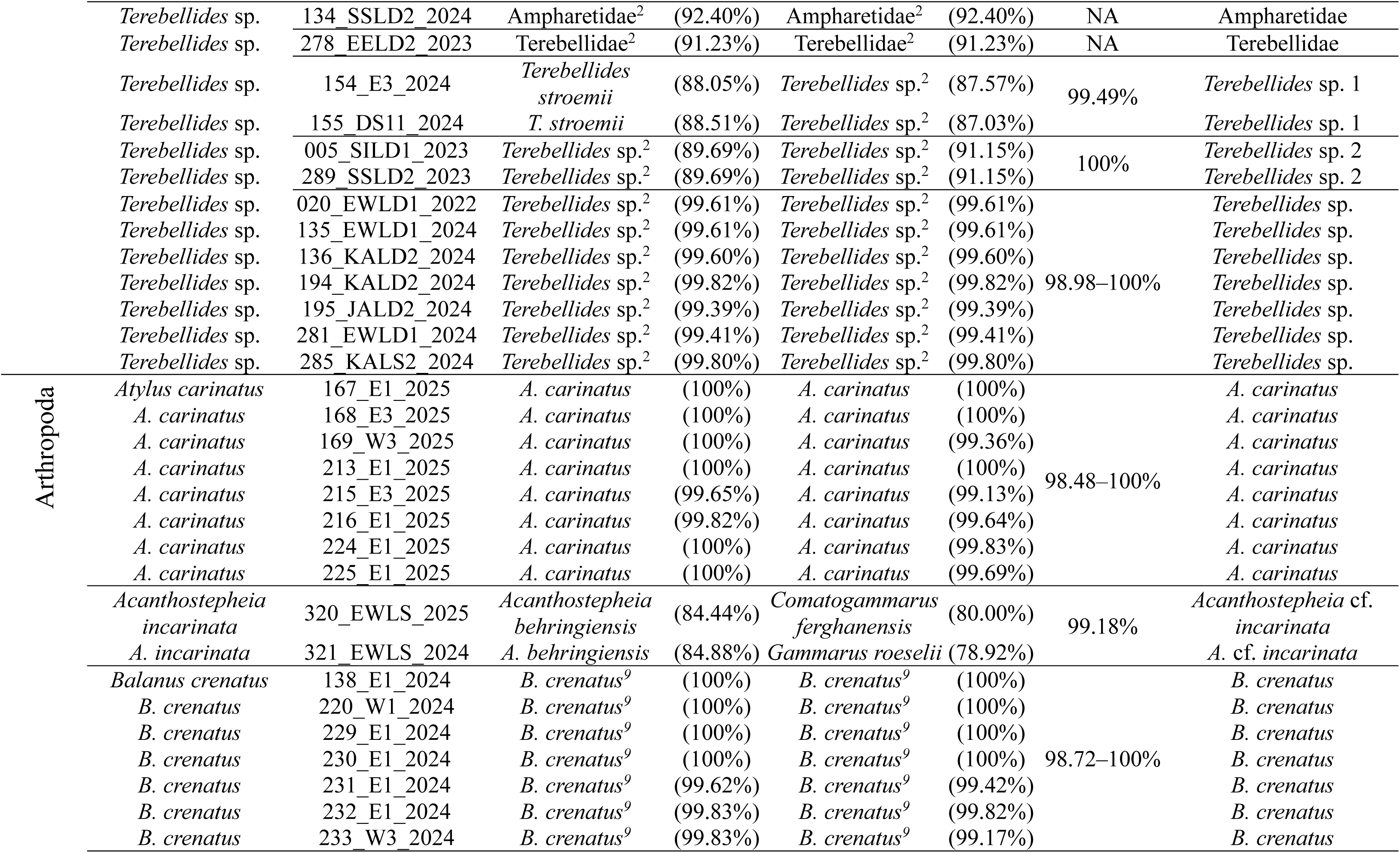
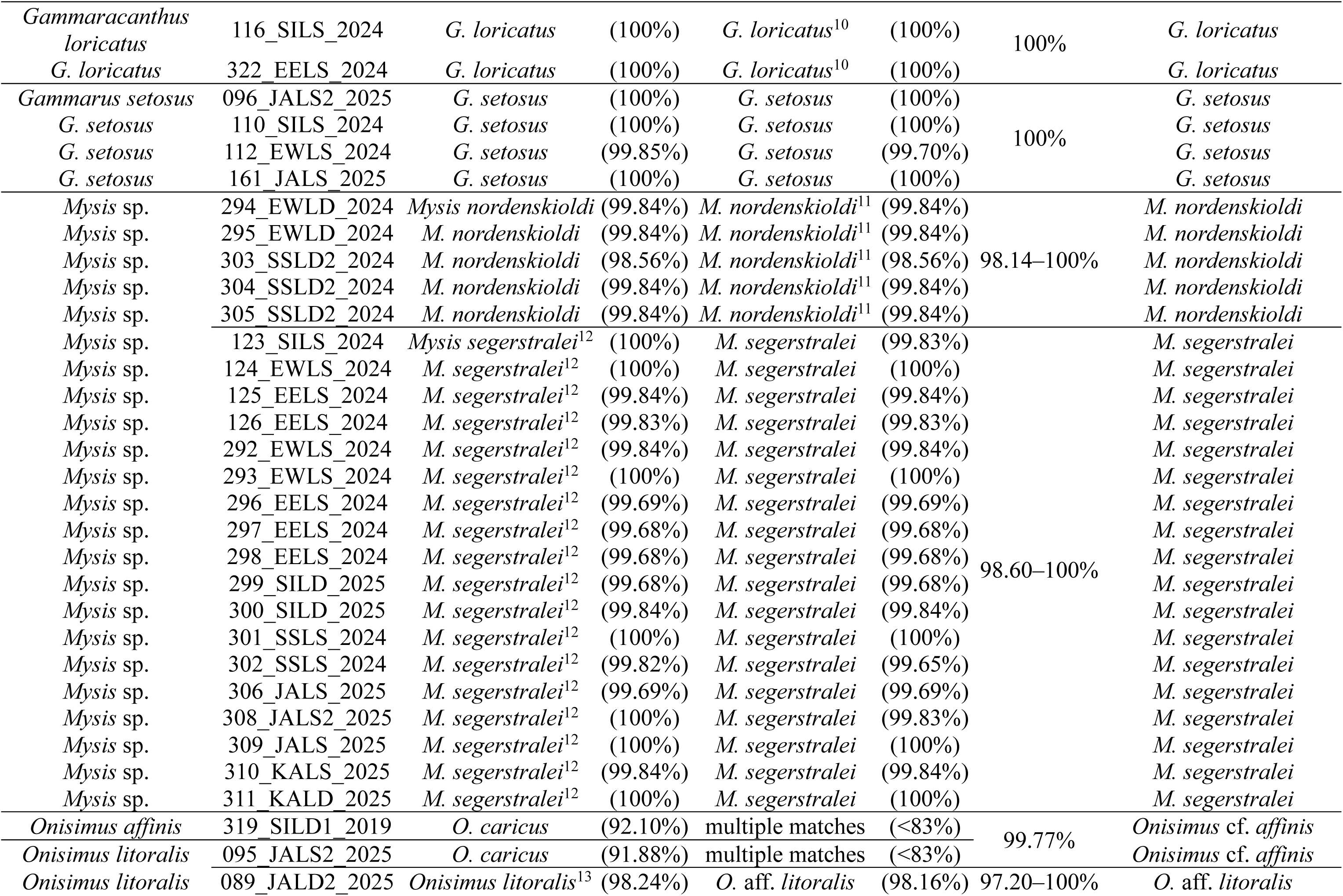
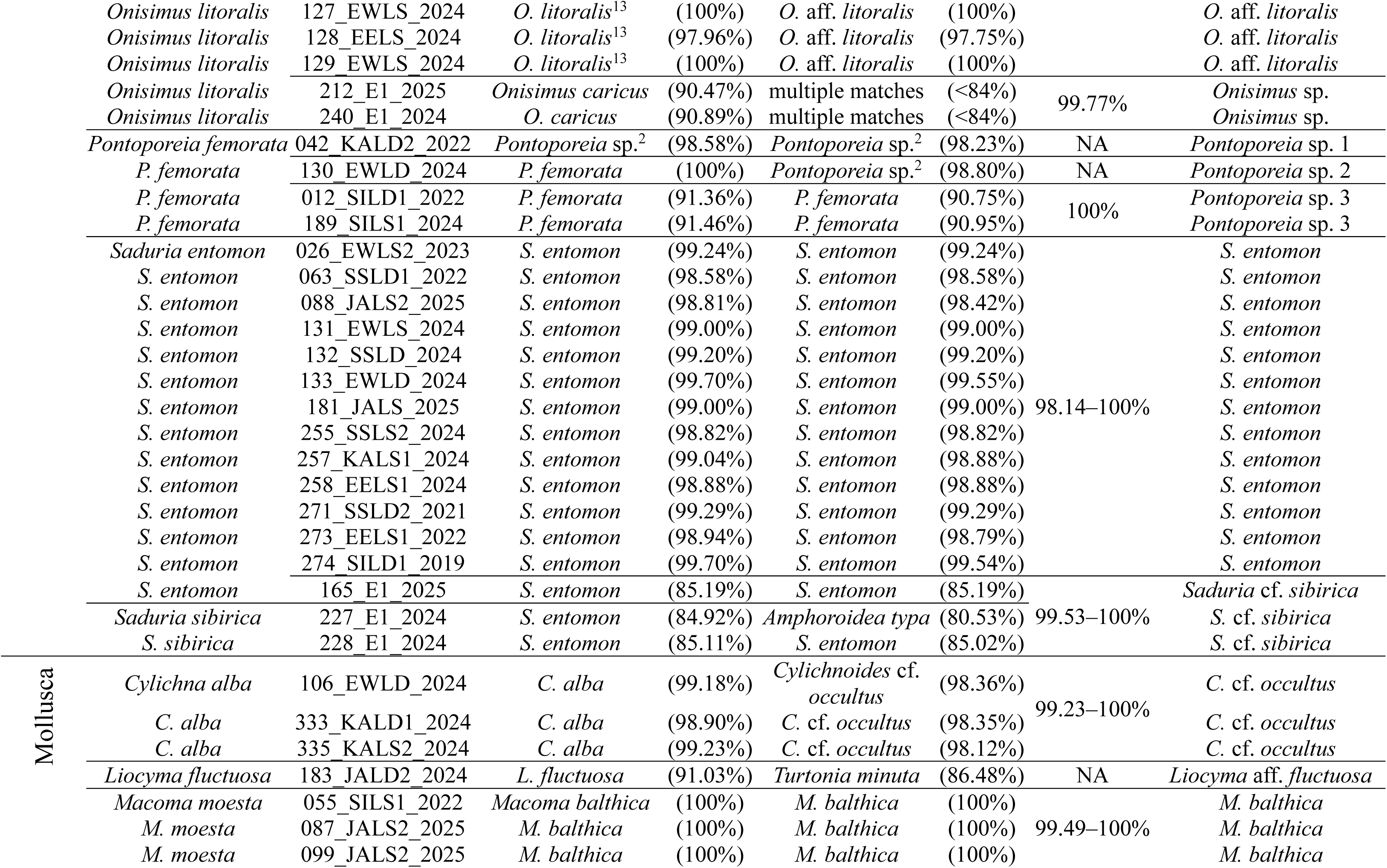
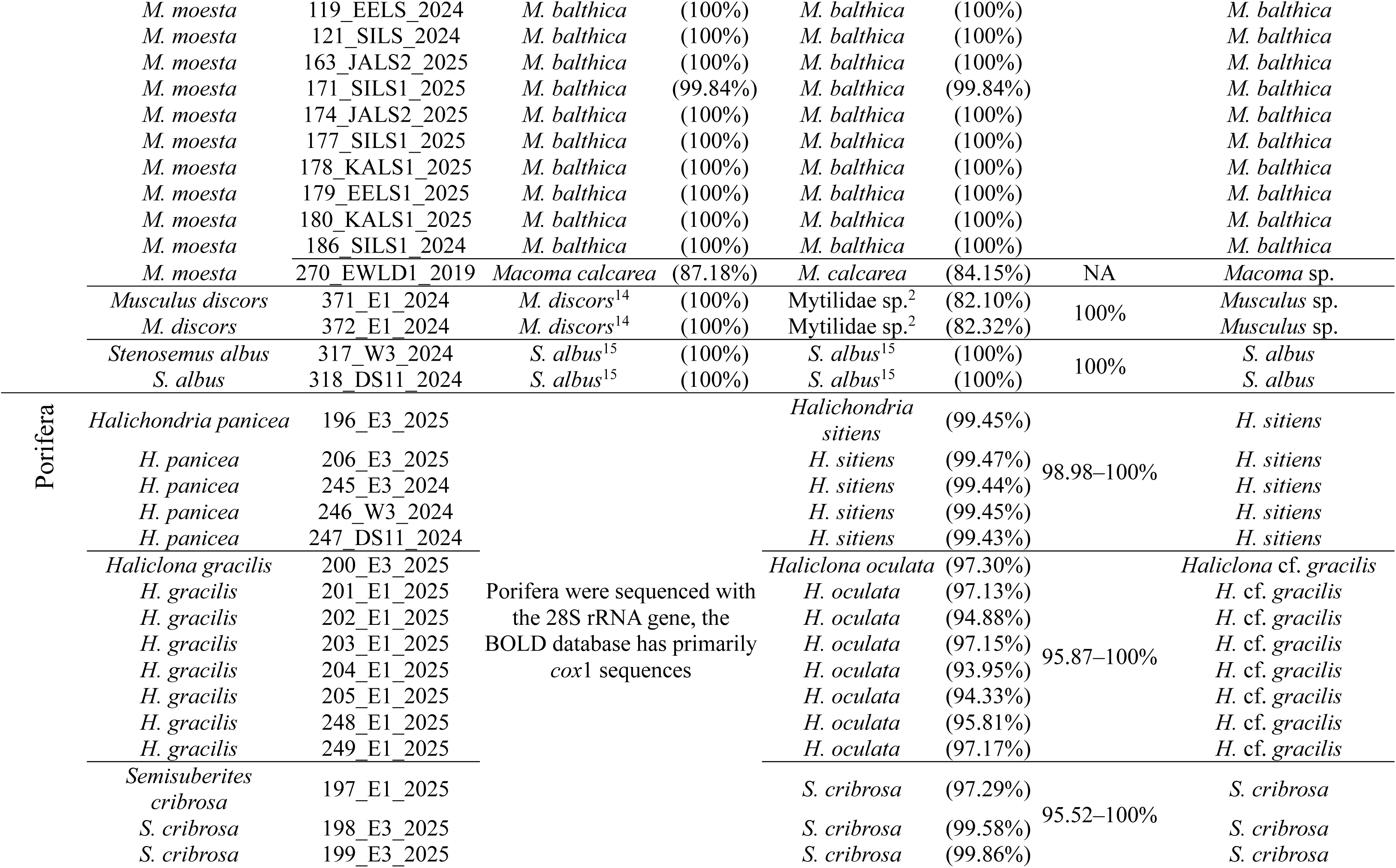
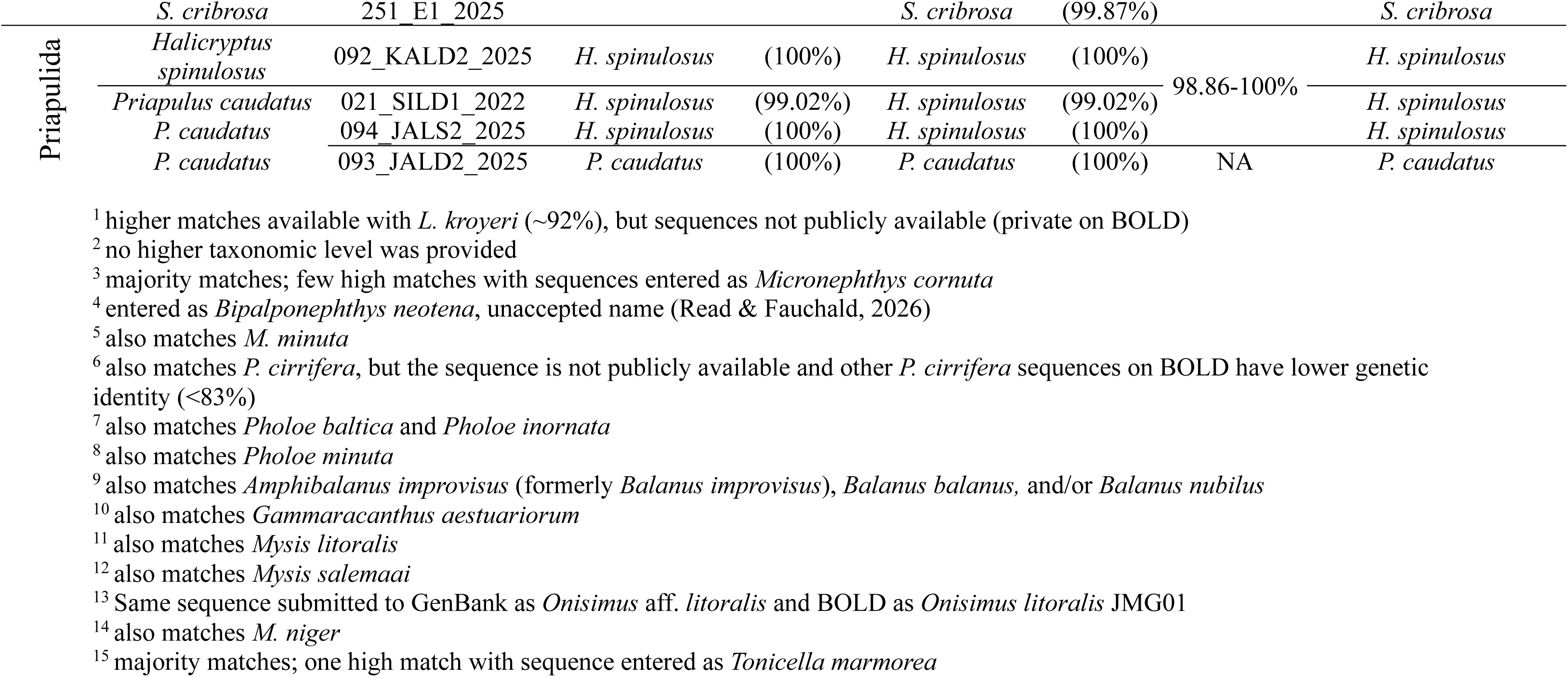
Comparison of morphological identification and genetic matches (BOLD and GenBank, except for Pori amplified samples. Intraspecific similarity of sequences from this study. Final taxonomy assignment based on m genetic identification.

All sequences for *C. polaris*, *N. aphroditoides* and *P. maculata* matched their morphological species identification (99.67-100%; Table 1). Whilst it was not possible to assign a species name to the two *Prionospio* sp. individuals based on our sequences, they had 100% genetic identity with *Prionospio* sp. 2.

The closest genetic matches for the individual morphologically identified as *E. nikolskyi* were species in the genera *Limnodrilus* and *Tubificoides* (<83% genetic identity). Publicly available sequences for *E. nikolskyi* had a genetic identity of only 78.95% with our sequences. In Figure 2, we only included species in *Limnodrilus* and *Tubificoides* for which the taxonomic placements are resolved (Kvist, Sarkar, & Erséus, 2010; Liu et al., 2017).

There are currently several sequences for *L. kroyeri* published in BOLD, which cluster separately (Figure 2) and not with our sequences for individuals identified as *L. kroyeri* based on morphology.

Of the sequences for the nine individuals morphologically identified as *M. minuta*, five matched published sequences for this species (genetic identity of 99.49-99.75%) without any discrepancies. Sequences for three other individuals matched published sequences for *M. neotena* (formerly *Bipalponephtys neotena*; genetic identity of 98.47-98.98%) in addition to sequences published as *M. minuta*.

In the genus *Pholoe,* multiple species are published under the same species names in genetic databases. Albeit not corrected on GenBank or BOLD, this was thoroughly reviewed by Meißner, Bick, & Götting (2017) and Meißner, Götting, & Nygren (2020) which we followed to inform taxonomy. Individuals in the genus *Pholoe* were not morphologically identified to species level. Of the five sequences we obtained, two matched *Pholoe assimilis* (99.24-99.49%) and three matched *Pholoe minuta* (98.72-99.75%).

We sequenced 13 individuals morphologically identified as *Terebellides* sp. which genetically represent five species (68.18-88.04% interspecific and 98.75-100% intraspecific genetic identity). The sequences of seven individuals for one species matched publicly available *Terebellides* sp. sequences (98.97-99.82% genetic identity, no higher taxonomic level provided) from several Arctic locations (Kattegat, Skagerrak, White Sea, and Canadian Arctic Archipelago). The sequences of four individuals belonging to two species have lower genetic identity with published *Terebellides* sequences (87.03-91.15%), but they still cluster within the genus. The sequence for 134_SSLD2_2024 had the highest genetic identity with a published sequence for Ampharetidae (highest taxonomic level provided, 92.40%). They clustered together and away from the *Terebellides*. The sequence for 278_EELD2_2023 had the highest genetic identity with a published sequence for Terebellidea (highest taxonomic level provided, 91.23%).

### Arthropoda

We successfully sequenced individuals identified as *Atylus carinatus* (n=8), *Acanthostepheia incarinata* (n=2), *Balanus crenatus* (n=7), *Gammaracanthus loricatus* (n=2), *Gammarus setosus* (n=4), *Mysis* sp. (n=23), *Onisimus affinis* (n=1), *Onisimus litoralis* (n=7), *Pontoporeia femorata* (n=4), *Saduria entomon* (n=14) and *Saduria sibirica* (n=2) based on morphology (Figure 3). Overall intraspecific genetic identity ranged from 97.20–100% within Arthropoda.

**Figure 3.**
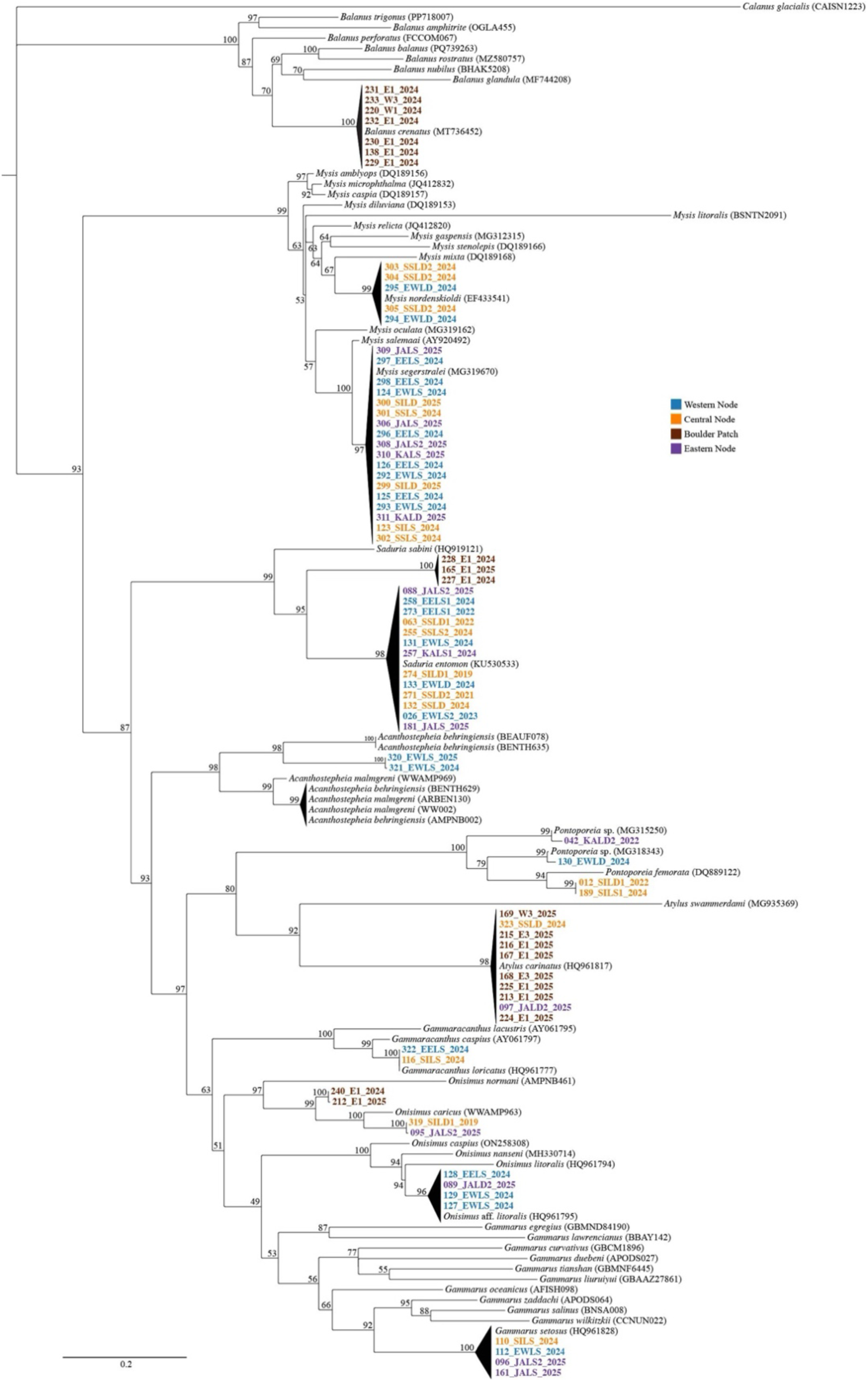
Maximum likelihood tree (*cox*1) for Arthropoda using the Transitional Model with empirical base frequencies, invariable sites and 4-tier Gamma distribution rates (TIM+F+I+G4). Values indicate bootstrap support. *Calanus glacialis* was used as the outgroup. Names in bold indicate sequences generated in this study with colors showing where they were collected from. All other sequences were sourced from the GenBank and BOLD databases (accession numbers provided).

All sequences for *A. carinatus*, *B. crenatus*, *G. loricatus*, *G. setosus* and all but one for *S. entomon* matched their morphological species identification (99.05-100%; Table 1). The sequence for the one *S. entomon* outlier clusters with sequences we generated for individuals morphologically identified as *S. sibirica* (99.53–100 % genetic identity). Since there are no sequences publicly available for *S. sibirica,* ours represent a novel contribution for these species. This is also the case for *A. incarinata*, for which the sequences here are the first published representatives.

There were some discrepancies for sequences published within *Balanus* and *Gammaracanthus*. Sequences generated for individuals identified morphologically as *B. crenatus* matched sequences published as various barnacle species: *B. crenatus*, *Amphibalanus improvisus* (formerly *Balanus improvisus*), *Balanus balanus,* and/or *Balanus nubilus*. Since most matches were for *B. crenatus* and independently sourced sequences for the other species did not cluster with our sequences, we kept *B. crenatus* as the species identification for these barnacles. Among multiple published sequences of *G. loricatus* only one was named *Gammaracanthus aestuariorum* and we, therefore, kept the former as the species name.

Based on molecular data we identified two species of individuals morphologically identified as *Mysis* sp.: *Mysis nordenskioldi* (formerly part of the *Mysis litoralis* species complex (Audzijonyte & Väinölä, 2007)) and *Mysis segerstralei* (formerly part of the *Mysis salemaai* species complex (Audzijonyte & Väinölä, 2006)) with 98.56-99.84% and 99.51-100% genetic identity respectively.

The sequences for individuals morphologically identified as *O. litoralis* belong to three different species based on molecular data (81.12–81.82% interspecific and 97.20–100% intraspecific genetic identity). Four of our sequences matched publicly available sequences for *Onisimus* aff. *litoralis* (97.75-100% genetic identity). Two sequences clustered together within the genus *Onisimus* and had a genetic identity of 90.47–90.89% with *Onisimus caricus*. The sequence for 319_SILD1_2019 matched the sequence we generated for the morphologically identified *O. affinis* (99.77% genetic identity). To our knowledge the only published sequence on BOLD for *O. affinis* is relatively short (170 base pairs) for an individual collected from Svalbard and does not match our sequence. Since we cannot verify this sequence without a linked publication, we are confident that our sequences represent the first entry for *O. affinis* from the Beaufort Sea.

The four individuals identified as *P. femorata* based on morphology represent three different species genetically (84.15–86.01% interspecific genetic). Available sequences on BOLD and GenBank are submitted as *P. femorata* and *Pontoporeia* sp. and are represented in two of the three genetic clusters. The sequences for 130_EWLD_2024 matched sequences on GenBank labeled *Pontoporeia* sp. whereas the sequence for the identical sample is labeled *P. femorata* on BOLD (sample ID NVAMP-0107 is the same across databases and BOLD links the GenBank sequence entry). These sequences cluster separately from other *P. femorata* entries on GenBank and BOLD which makes it impossible to resolve their species identification.

Sequences for two of our individuals cluster together and did not have any close matches (90.75–91.46% genetic identity with *P. femorata* public sequences).

### Mollusca

We successfully sequenced individuals identified as *Cylichna alba* (n=3), *Liocyma fluctuosa* (n=1), *Macoma moesta* (n=14), *Musculus discors* (n=2), and *Stenosemus albus* (n=2) based on morphology (Figure 4). Overall intraspecific genetic identity ranged from 99.23–100% within Mollusca.

**Figure 4.**
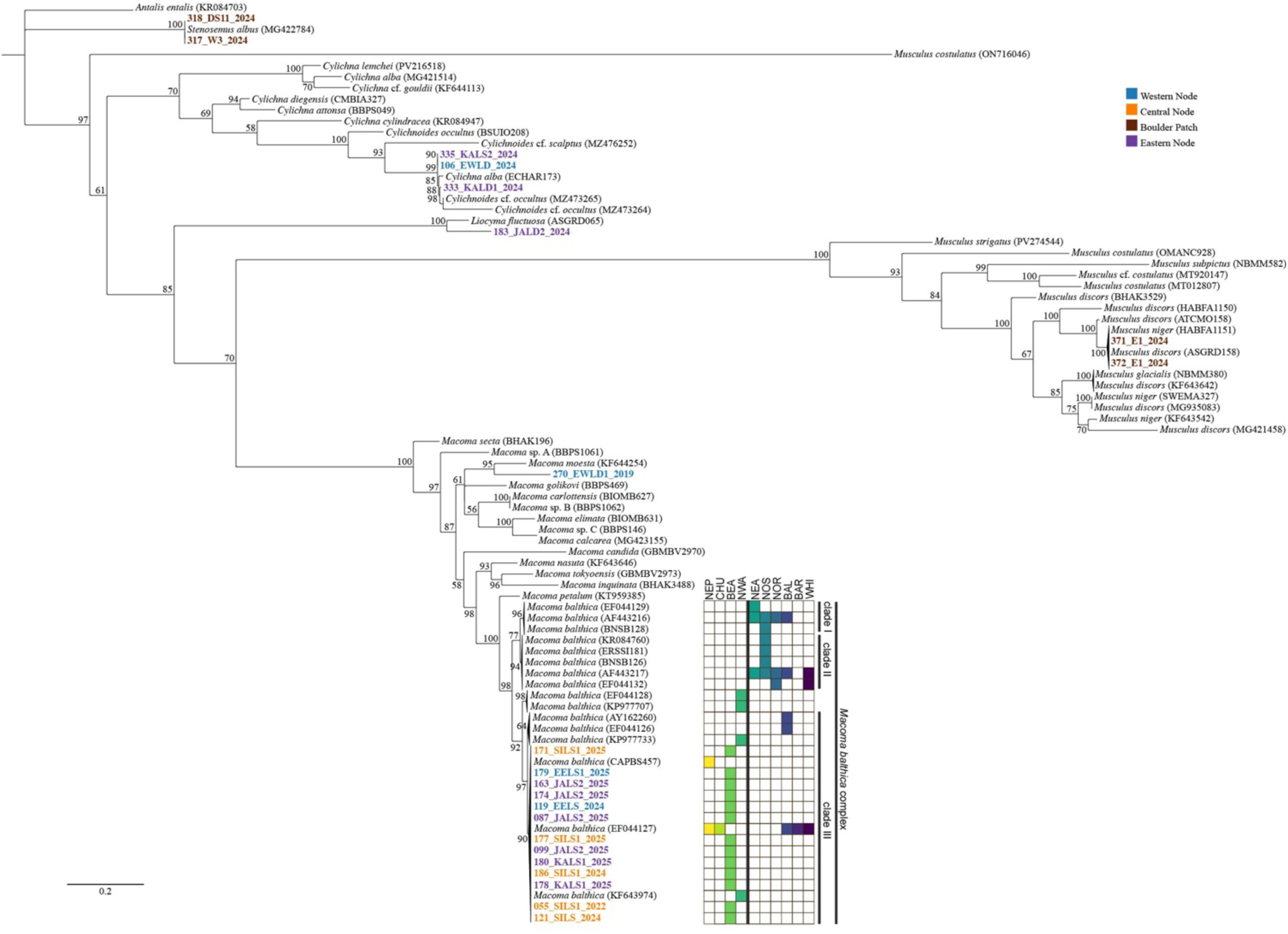
Maximum likelihood tree (*cox*1) for Mollusca using the Transitional Model with empirical base freque Gamma distribution rates (TIM+F+G4). Values indicate bootstrap support. *Antalis entalis* was used as the outgro indicate sequences generated in this study with colors showing where they were collected from. All other sequen from the GenBank and BOLD databases (accession numbers provided). Heatmap shows where each sequence w – Northeast Pacific, CHU – Chuckchi Sea, BEA – Beaufort Sea, NWA – Northwest Atlantic, NEA – Northeast A Sea, NOR – Norwegian Sea, BAL – Baltic Sea, BAR – Barents Sea, and WHI – White Sea. The vertical line sep adjacent to North America from those adjacent to Europe.

The two sequences for *S. albus* matched their morphological identification (Table 1). The three sequences for individuals identified morphologically as *C. alba* matched BOLD entries for *C. alba* (98.90–99.23% genetic identity) and GenBank entries for *Cylichnoides* cf. *occultus* on GenBank (98.12-98.36% genetic identity). Since our sequences cluster with other sequences in the genus *Cylichnoides* (Figure 4), we accept *C.* cf. *occultus* as the species name. The sequence for the individual identified as *L. fluctuosa* based on morphology matched one sequence in the BOLD database for the same species with 91.03% genetic identity. Our sequences for individuals identified as *M. discors* based on morphology matched *M. discors* (100% genetic identity) and *Musculus niger* (99.35–99.45% genetic identity) sequences in BOLD. In fact, many sequences submitted under these two species names cluster separately from each other (Figure 3).

The sequence for one individual of *M. moesta* (270_EWLD1_2019) based on morphology clustered separately from the other individuals sequenced (genetic identity of 84.15-87.18% with *Macoma calcarea*). All other sequences for the morphologically identified *M. moesta* matched published sequences for *Macoma balthica* (99.84-100% genetic identity). When analyzing these sequences, we also included sequences from numerous studies which investigated the trans-Arctic dispersal patterns of *M. balthica* (Barco, Raupach, Laakmann, Neumann, & Knebelsberger, 2016; Layton, Martel, & Hebert, 2016; Luttikhuizen, Drent, & Baker, 2003; Nikula, Strelkov, & Väinölä, 2007). Our sequences were subsequently assigned to *M. balthica* clade III based on (Luttikhuizen et al., 2003) which coincides with sequences previously classified as *Macoma balthica balthica*. *M. balthica* clades I and II (*Macoma balthica rubra*) have thus far been found in the Northeast Atlantic, North Sea, Norwegian Sea, Baltic Sea, Barents Sea, and White Sea, whereas clade III has been found in most of those locations and in the Northeast Pacific, Chuckchi Sea, Beaufort Sea, and Northwest Atlantic (Figure 4).

### Porifera

We successfully sequenced an 800 base pair region of the 28S rRNA gene for individuals morphologically identified as *Halichondria panicea* (n=5), *Haliclona gracilis* (n=8), and *Semisuberites cribrosa* (n=4; Figure 5). Overall intraspecific genetic identity ranged from 95.52–100% within Porifera which was higher than expected due to ambiguities in several sequences that could not be resolved. All the ambiguities were found in the same part of the sequences. When we removed this portion, leaving 640 base pairs, intraspecific genetic identity ranged from 98.31–100%.

**Figure 5.**
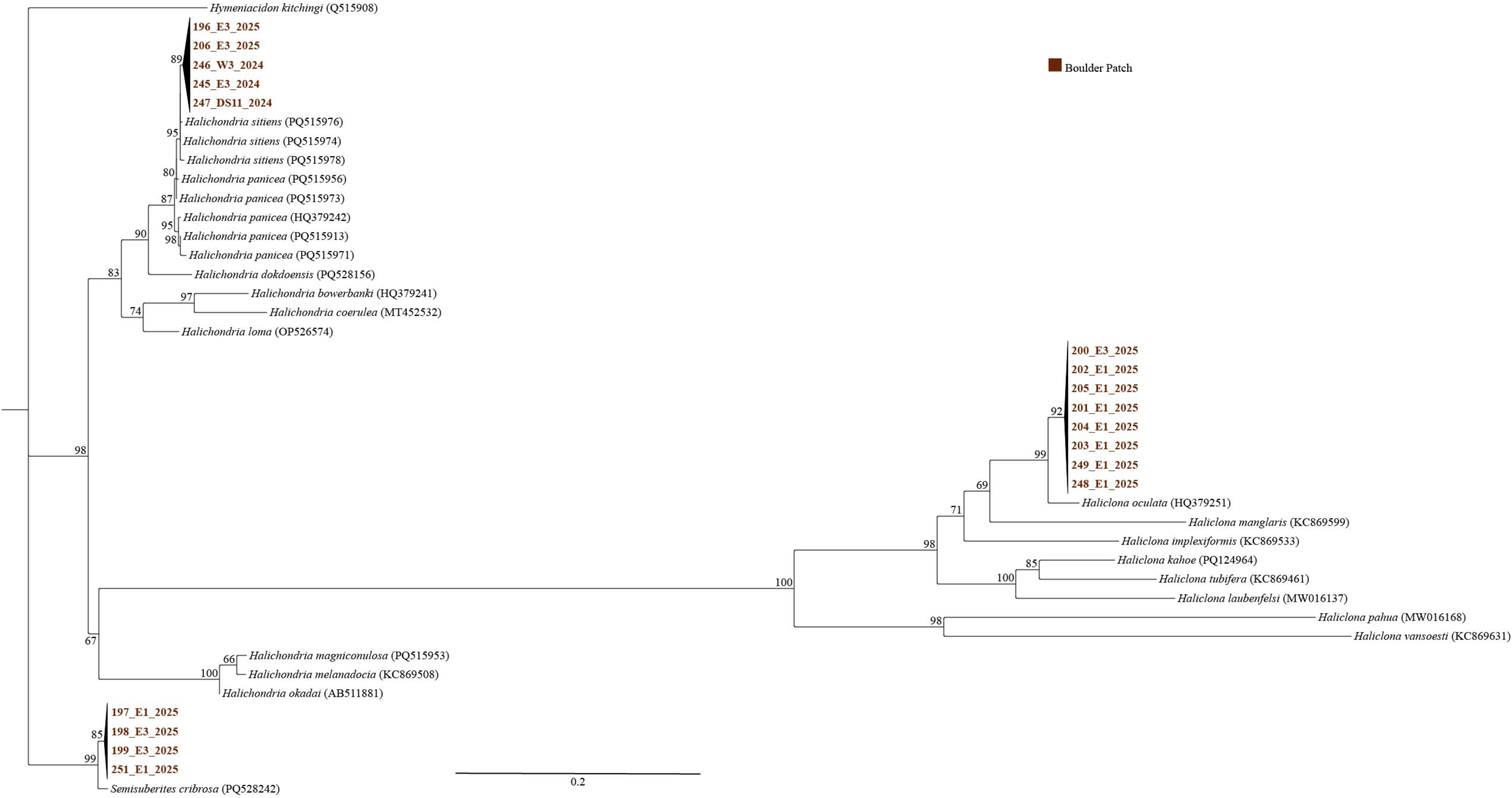
Maximum likelihood tree (28S rRNA gene) for Porifera using the Transitional Model 3 with empirical 4-tier Gamma distribution rates (TIM3+F+G4). Values indicate bootstrap support. *Hymeniacidon kitchingi* was u Names in bold indicate sequences generated in this study with colors showing where they were collected from. A were sourced from the GenBank and BOLD databases (accession numbers provided).

All sequences for individuals identified as *S. cribrosa* based on morphology matched database entries for the same species (97.29-99.87% genetic identity, 99.62–99.81% genetic identity for trimmed sequences; Table 1). The sequences for the morphologically identified *H. panicea* matched sequences for *Halichondria sitiens* (99.43-99.47% genetic identity, no ambiguities in the sequences) followed by matches with sequences for *H. panicea* (99.15-99.22% genetic identity). These two species are very closely related and mostly distinguished by spicule length and surface papillae (Turner, Morrow, Picton, Goodwin, & Thacker, 2024) which was not measurable as part of this study. Our sequences cluster with other *H. sitiens* sequences (Figure 5, Dr. Thomas Turner *pers. comm.*) and we therefore believe that this is the species collected from the Boulder Patch. The sequences for individuals identified morphologically as *H. gracilis* are closest to public sequences for *Haliclona oculata* (93.95–97.30% genetic identity; 97.57–97.76% genetic identity for trimmed sequences). Since genetic identity is below the 98% species threshold and there are no sequences for *H. gracilis* in GenBank (BOLD was not consulted since it is specific to *cox*1 sequences), this likely represents the first contribution for this species.

### Priapulida

We obtained sequences for individuals identified as *Halicryptus spinulosus* (n=1) and *Priapulus caudatus* (n=3) based on morphology (Figure 6). Overall intraspecific genetic identity ranged from 98.86–100% within Priapulida. Three of the Priapulida sequences matched sequences for *H. spinulosus* (99.02-100% genetic identity; Table 1). The sequence for one individual morphologically identified as *P. caudatus* matched sequences for the same species (100% genetic identity).

**Figure 6.**
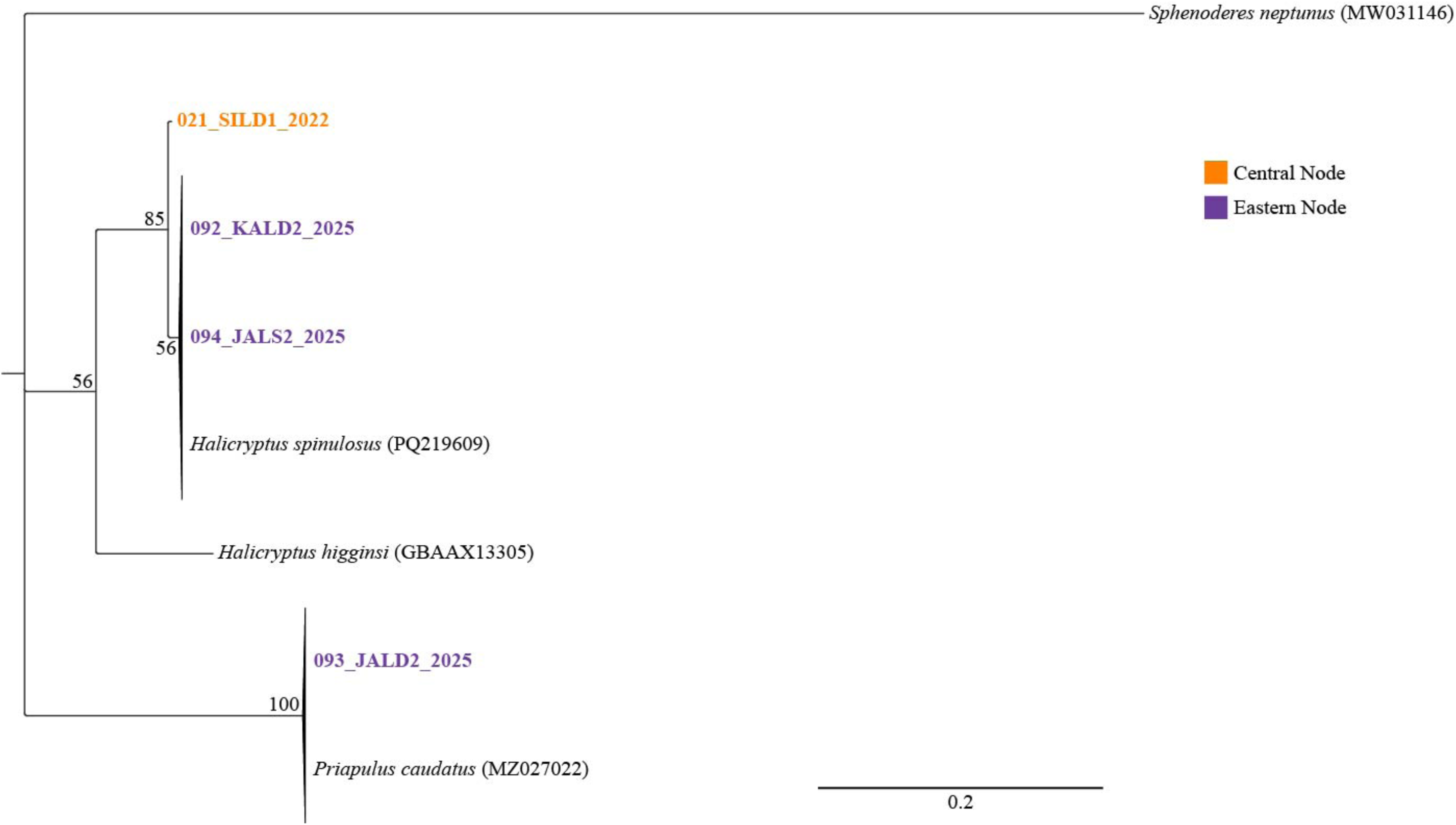
Maximum likelihood tree (*cox*1) for Priapulida using the Kimura 3-parameter model with empirical ba tier Gamma distribution rates (K3Pu+F+G4). Values indicate bootstrap support. *Sphenoderes neptunus* was used Names in bold indicate sequences generated in this study with colors showing where they were collected from. A were sourced from the GenBank and BOLD databases (accession numbers provided).

## Discussion

Overall, sequencing revealed a higher species diversity compared to morphological identification (morphospecies): 15 compared to nine morphospecies for Annelida, 15 compared to 12 morphospecies for Arthropoda, and six compared to five morphospecies for Mollusca.

Whilst the number of species for Porifera and Priapulida were the same between methods, molecular analyses did not always agree with the morphospecies. Overall, we could not match all putative cryptic species we identified using molecular methods with sequences in the databases since this region of the world and some of the taxa remain understudied. Multiple genetic markers need to be used to confirm our findings and accurately unravel the species diversity, especially if they are truly cryptic and no distinguishing morphological features are present.

We also generated sequences for four morphospecies that do not have any known published entries in databases (*A. incarinata*, *H. gracilis*, *O. affinis*, and *S. sibirica*). Until it is possible to sequence type specimens for these morphospecies, we can use our sequences as preliminary references. The number of genetically described species is much higher compared to morphospecies for Annelida, which is partially due to the difficulty of identifying some taxa (here *Pholoe* and *Terebellides*) beyond genus level. Resources such as time and expertise in all taxonomic groups are limiting factors when broad biodiversity studies are being conducted, which can also lead to misidentification of some individuals. Our research emphasizes that morphological species identification alone underestimates biodiversity and needs to be combined with molecular analyses. This work is crucial to build reliable genetic databases with representation of common taxa in the ecosystems being studied (Pappalardo et al., 2025), so we can detect future changes in species diversity.

### Cryptic diversity

Concepts for distinguishing among species are widely discussed by taxonomists, and here we are identifying potential cryptic species complexes based on our molecular investigations. We used generalized genetic identity thresholds commonly applied across taxonomic groups such as 98% for species assignment (deWaard et al., 2019; Hebert, Ratnasingham, & deWaard, 2003) in addition to phylogenetic tree topology. However, inter- and intraspecific genetic differentiation can vary depending on the genetic marker used as well as among phyla (Huang, Meier, Todd, & Chou, 2008) and families (Lefébure, Douady, Gouy, & Gibert, 2006; Tempestini, Rysgaard, & Dufresne, 2018). The threshold is usually determined by the barcoding gap where interspecific divergence consistently exceeds intraspecific variability. All but one of our samples for *Macoma* matched sequences of the *M. b. balthica* lineage (clade III in Figure 4) in the well-studied *M. balthica* complex (99.49–100% intraspecific genetic identity, 81.38–81.89% interspecific genetic identity). Keeping the one outlier as *M. balthica* instead of accepting it as a cryptic species would artificially increase the range of intraspecific genetic identity *M. b. balthica*. However, it is noteworthy that the *M. b. rubra* lineage (clades I and II in Figure 4) is capable of doubly uniparental inheritance where male and female mitochondrial DNA are passed to male offspring and can be highly divergent (Le Cam et al., 2025). Our outlier *Macoma* sequence did have several discrepancies (unclear peaks or disagreement between forward and reverse sequence). In the future, it might help to investigate intra-individual variation for this and other individuals to determine whether there are highly divergent versions of the *cox*1 gene present. Thus, genetic sequencing could lead to overestimation of biodiversity (Martínez, Harms, Abele, & Held, 2023). Whether or not this is the case here needs further investigation.

Whilst combining multiple cryptic species under one species name may inflate intraspecific genetic differentiation, having gaps in species sampling when resolving phylogenetic relationships may overestimate interspecific differentiation values. Since *Pholoe* and *Terebellides* were only identified to genera, finding multiple species within each was expected. However, we did not anticipate that four individuals with the assigned morphospecies *Pontoporeia femorata* would genetically resolve into three different genetic clusters (83.44-85.93% genetic identity). To date, there is only one accepted morphospecies within the genus *Pontoporeia* (Horton et al. 2026). Combining public records and our novel *Pontoporeia* spp. suggests that there are at least four cryptic species in this genus. Basing species description on morphology alone only works when distinguishing features are known.

This is also the case in the annelid *Micronephthys* where we discovered *M. neotena* in addition to the morphospecies *M. minuta*. Individuals in the genus *Micronephthys* have been known to be misidentified, and additional cryptic diversity is suspected (Dnestrovskaya & Jirkov, 2010) which can then lead to incorrectly submitted genetic sequences (discussed later). In Northern Europe and the Arctic, *M. neotena* is generally found in slightly warmer conditions than *M. minuta*, but both have been found to co-occur (Dnestrovskaya & Jirkov, 2010). However, when they do *M. neotena* appears to outcompete *M. minuta* (Dnestrovskaya & Jirkov, 2010) which may explain why three of our four *Micronephthys* in Elson lagoon belonged to *M. neotena.* This species was only found in Elson lagoon which is closer to the Pacific and boreal habitats than the other sites. Whether or not this reflects a more recent arrival from the Pacific due to global warming requires barcoding of older samples from this lagoon. In addition, there is evidence that a minimum sample size of 100 individuals of the annelid genus *Terebellides* is required to accurately determine species diversity within a biogeographic region (Nygren et al., 2018). This suggests that larger sample sizes are necessary to uncover the full range of cryptic species across taxa in the lagoons of the Beaufort Sea.

In our study, cryptic species include those for which no visible distinguishing features have been identified yet. In other cases, specific expertise may be required to tell species apart morphologically as is the case with the sponges *H. sitiens* and *H. panicea* in which spicule length is a distinguishing feature (Turner et al., 2024). This is common in sponges where species have been delimited by microscopic features such as spicule morphology and molecular sequences with as little as single base pair differences in the D3–D5 and D6–D8 rRNA gene regions (Dinn, Edinger, & Leys, 2019). Thus, molecular or morphological methods alone are not always sufficient to delineate species.

If cryptic species are geographically isolated and they are not recognized, this can lead to regional endemism being underestimated. The notion that the Arctic is characterized by low endemism has been revised, especially since the increased use of molecular techniques. For example, around 55% of macroalgal morphospecies described from the Boulder Patch were renamed after molecular analyses (Bringloe, Dunton, & Saunders, 2017). Extreme environments, such as polar regions, have been postulated to support morphological stasis during speciation, leading to an increased number of cryptic species (Bickford et al., 2007). In polar polychaetes 25-50% of morphospecies examined contained more than one cryptic species (Brasier et al., 2016; Carr, Hardy, Brown, Macdonald, & Hebert, 2011). Without full description of cryptic species diversity within biogeographic regions, we will not be able to accurately assess future changes such as the arrival of non-native or disappearance of other species. In addition, cryptic species within a morphospecies complex may be more limited in their geographic distribution, which in turn implies a smaller population size and a higher chance of extinction (Bickford et al., 2007).

Cryptic diversity can be facilitated by niche differentiation. The Boulder Patch in the Beaufort Sea constitutes a highly unique ecological oasis due to its hard substratum in an otherwise soft sediment dominated ecosystem. It is therefore not surprising that it supports a very different invertebrate community assemblage (Dunton et al., 1982). Whilst some species were found in soft and hard substratum habitats, others appear limited to one or the other, bearing in mind that due to resource limitations we focused mainly on the most common taxa or species for which no genetic sequences are available. The most unbiased collections were *Pholoe* spp. and *Terebellides* spp. since morphological identification was limited to genus level. In both genera, the species we identified using molecular methods showed strong habitat preference, where *P. assimilis* and one unidentified *Terebellides* sp. were restricted to the Boulder Patch. Similarly, molecular analysis of the common morphospecies *O. litoralis* revealed cryptic diversity, with an unidentified species found in the Boulder Patch in addition to *O.* aff. *litoralis* found at soft bottom sites. Instituting more in-depth molecular studies in the Boulder Patch will help discover its true biodiversity profile.

### Trans-arctic connectivity

Many Arctic species have temperate counterparts in the adjacent oceans which are either the same species with tolerances for broad environmental factors (e.g. temperature) or closely related species where one is endemic to the Arctic (Bringloe et al., 2020; Morozov, Sabirov, & Anisimova, 2021; Morozov, Sabirov, & Zimina, 2018; Tempestini et al., 2018). Whether the Atlantic or Pacific has greater contributions to Arctic biodiversity is still debated. More amphipod species found in the Canadian Arctic are shared with the Atlantic than the Pacific (Tempestini et al., 2018). However, marine forests in the east Canadian Arctic are a mix of Pacific and Atlantic origin whereas the west Arctic is more similar to the Bering Sea in the northern Pacific (Bringloe et al., 2020). Unraveling these patterns of trans-Arctic connectivity is complicated by misidentification of species with only minute morphological differences (Morozov et al., 2021) which further supports the importance of including molecular data into these analyses. Additionally, discontinuous distributions of some species suggest past invasions and displacement in some areas, driven by local ocean current patterns supporting the arrival of competitors and/or removal of larvae (Kudryavtseva et al., 2023). Benthic taxa with planktonic life cycle stages, especially if their duration is longer, may be more prone to trans-Arctic dispersal (Hardy et al., 2011). The *M. balthica* complex is among the best studied trans-Arctic species (Barco et al., 2016; Layton et al., 2016; Luttikhuizen et al., 2003; Nikula et al., 2007). *M. b. rubra* has been recorded from the Northeast Atlantic and adjacent seas, whereas *M. b. balthica* has been found from the Northeast Pacific through the Arctic to the White Sea. Our data from the Beaufort Sea add a connection between the existing molecularly supported reports from the Chukchi Sea and the Northwest Atlantic (Layton et al., 2016).

Since the opening of the Bering Strait, it has been possible for species to reach the Atlantic from the Pacific through the Arctic (Bringloe & Saunders, 2019). In the Beaufort Sea one current runs towards the Canadian Arctic along the coastline whereas the Beaufort Gyre moves water clockwise and therefore in the opposite direction towards the East Siberian Sea. From there the Transpolar Current, which is the strongest Arctic Ocean current, transports water straight across the Arctic towards the Atlantic. Hypothetically, these currents provide multiple avenues for organisms, especially planktonic larval stages, to travel across the Arctic. Several of the polychaete species we sequenced from the Beaufort coastline have broader distributions in this polar region. *Pholoe minuta*, for example, is among the most widely distributed pan-Arctic polychaete species found on the Arctic shelves (Piepenburg et al., 2011). Most of our *Terebellides* sequences matched those of an undescribed species within this genus assigned to clade 4. This clade has been found in the Canadian Arctic Ocean (Resolute Bay), North Sea (Skagerrak), Kattegat (strait connecting the North and Baltic Sea), and the White Sea where it appears to be more commonly found in relatively shallow (<20 m) and enclosed areas where they are able to withstand salinity fluctuations (Carr et al., 2011; Kudryavtseva et al., 2023; Nygren et al., 2018) - conditions which are comparable to those at our coastal Beaufort Sea sites. Other species, such as *Prionospio* sp. 2 have been recorded in the Laptev Sea (Hektoen, Bakken, Ekrem, Radashevsky, & Dunshea, 2024). Many of the species we identified claim circumpolar distribution. However, most of them lack molecular studies to determine whether they are truly the same species or represent more examples of cryptic species. Here we confirm that many species which have been sequenced in other areas of the Arctic are also found in coastal areas of the Beaufort Sea which supports its potential importance as a steppingstone for species to disperse from the Pacific to the Atlantic Ocean.

### Genetic databases

Our ability to match the taxonomy of individuals based on molecular methods without morphological identification (e.g. via metabarcoding), is limited by the accuracy of public genetic databases. As we expand our knowledge of cryptic species and resolve their genetic relationships, database entries for previously published sequences should be confirmed and revised, if warranted. In addition, having sequences published in different databases can lead to further uncertainties for the correct taxonomy especially when no publication is linked to the entries. In the case of *Pontoporeia*, the same sequence for the same sample was submitted as *P. femorata* to BOLD and *Pontoporeia* sp. to GenBank. There are additional entries under *P. femorata* which only have an 85% genetic identity with the former *P. femorata* entries. Since the average intrageneric divergence for *cox*1 in Crustaceans is 17.16% (Costa et al., 2007), sequences submitted as *P. femorata* likely represent another cryptic species complex which warrants further investigation.

For the phylum Annelida, we identified discrepancies for multiple species. Within *Micronephthys*, sequences belonging to the same species were submitted under *M. minuta* and *M. neotena*. Other sequences, which only had a ∼94.5% genetic identity with these sequences, were all published under *M. minuta*. Two independent publications describing sequences published as *M. neotena* identified their specimens morphologically prior to sequencing (Carr et al., 2011; Huč, Hiley, McCowin, & Rouse, 2024). We therefore followed these studies for species assignment of *M. neotena* based on barcoding. Since we also were unable to distinguish between the two species based on morphology, it is not surprising that sequences have been published under both names. *Pholoe* spp. and *Terebellides* spp. are also notoriously difficult to distinguish for those who are not specialized in the genera. Some of the incorrectly published sequences for *Pholoe* spp. have been thoroughly reviewed (Meißner et al., 2017, 2020) which we relied upon, as records in GenBank or BOLD have not been revised. For the *Terebellides*, Figure 2 includes two *T. stroemii* sequences, since it was not possible to identify the true representative for this species. However, because none of our sequences matched *T. stroemii*, we did not thoroughly investigate this discrepancy. Similarly, public *E. nikolskyi* sequences did not match barcoding data for our morphologically identified *E. nikolskyi*. The specimens for the already published sequences originated from freshwater lakes in Japan (Ohtaka & Martin, 2011) and Russia (Martin, Sonet, Smitz, & Backeljau, 2019). Even though many of our sites are in lagoons with significant freshwater input during certain times of the year, conditions are very different and include large environmental fluctuations in salinity. Since we could not compare our individuals morphologically or genetically to type specimens, we refer to our samples as *E.* aff. *nikolskyi.* Including open nomenclature qualifiers is helpful when available morphological identification does not agree with molecular data and the discrepancies have not been resolved in the literature.

When published sequences are not linked to scientific papers, it is much more difficult to verify which entry is correct. Our sequences for the morphologically identified *Musculus discors* matched sequences for both *M. discors* and *M. niger* on BOLD. However, without a linked publication it is unclear whether the authors for the different entries were aware of the discrepancy and what basis they used for their final identification. Without comparisons to type specimens or other reliable quality control steps, it should be mandatory to include qualifiers such as “cf.” or “aff.” to signify caution. Besides simply comparing sequences to public databases, generating phylogenetic trees can also resolve some discrepancies. Our sequences for *C. alba* matched entries under the same species name on BOLD but entries for *C.* cf. *occultus* on GenBank. Sequences for species in the family Eoscaphandridae (e.g. *C.* cf. *occultus*) cluster separately from those in the family Cylichnidae (e.g. *C. alba*; (Siegwald, Oskars, Kano, & Malaquias, 2022)). Since our sequences clustered clearly with *C.* cf. *occultus* and other species within this genus, we revised our morphological identification based on the molecular data. Taking multiple lines of evidence into account when assigning species names to molecular data is crucial for database integrity.

## Conclusion

Although undoubtedly some progress has been made since Piepenburg et al. (2011), their work suggests that 20–33% of Mollusca-Arthropoda-Echinodermata and 10–25% of Annelida macro- and megabenthic species are still unknown for the Arctic shelves. The history of the area promotes cryptic species diversity through times of glaciation where species retreated into refugia, followed by reopening of areas with reintroductions from southern areas. Almost 19% of our sequenced individuals belonged to species for which no sequencing data are currently available. This may be an underestimation of unrecorded biodiversity due to low sample size and non-random selection of morphospecies in this study. Nygren et al. (2018), for example, found that 33-50% of *Terebellides* in their study belonged to previously undescribed species. Taking inventory of the current biodiversity is crucial for detecting future changes. We have shown that it is necessary to include molecular in addition to morphological identification of specimens due to high cryptic diversity in Arctic ecosystems. Additionally, molecular data will help to further unravel the trans-Arctic dispersal pathways that have led to the current species distributions.

This work emphasizes that substantial effort is required to describe and name the cryptic species found in the Beaufort Sea as well as a thorough revision of public databases. This will require an increase in sample size, detailed morphological description, and the sequencing of multiple barcoding genes. Such data are vital for metabarcoding studies increasingly used to evaluate biodiversity and monitor changes in species composition including the arrival of non-native species. Initiatives such as the one led by the Icelandic Institute of Natural History where they process archived collections are a great example of how to build a reliable foundation for future biodiversity assessments (Gudmundsson, 2023). Their digital database based on morphospecies along with physical specimens provides a strong foundation for linking morphological and genetic species identifications. Worldwide efforts are being undertaken to build reliable, comprehensive barcoding databases to assess current biodiversity and monitor future changes. The international Barcode of Life (iBOL) initiative for example has multiple ongoing projects in Costa Rica (BioAlfa), the United Kingdom (BIOSCAN-UK), Europe (BIOSCAN Europe), and Canada (Arctic BIOSCAN). Molecular approaches help to mitigate the vast underrepresentation of marine non-vertebrate species in addition to terrestrial insects, fungi and soil invertebrates (Hobern, 2020). Moving forward, we will use our results to revise MIDORI2 (Leray et al., 2022), a reference database used for taxonomic assignment of *cox*1 metabarcoding data, generating an Arctic-specific version (Arctic MIDORI2; Heiser, *in prep.*) that will allow for more accurate species assemblage descriptions.

## Acknowledgments

We acknowledge that much of our research occurs on the traditional land that is owned and managed by the communities of Utqiaġvik, Nuiqsit, and Kaktovik. Our field sites overlap with Iñupiaq hunting and fishing grounds and sites of cultural and heritage significance. We commit to acknowledge, respect, and make our relationship visible to these people.

We would like to thank all members of the BLE LTER and Boulder Patch field teams who have collected, identified, and preserved the specimens over the years. This work would have not been possible without the help from Sharon Magnuson and Laurel Weeks from the TAMUCC Genomics Core Lab. We appreciate Drs. Mark Lever’s and Thomas Turner’s advice during the data analysis. We are thankful the taxonomic expertise contributions from Leslie Harris (polychaetes), Nora Foster (molluscs) and Ken Coyle (crustaceans).

## Data Availability Statement

Molecular data generated have been deposited in BOLD and GenBank (see results and Supplementary information for details).

## Supporting Information

**Supplemental Table 1.**
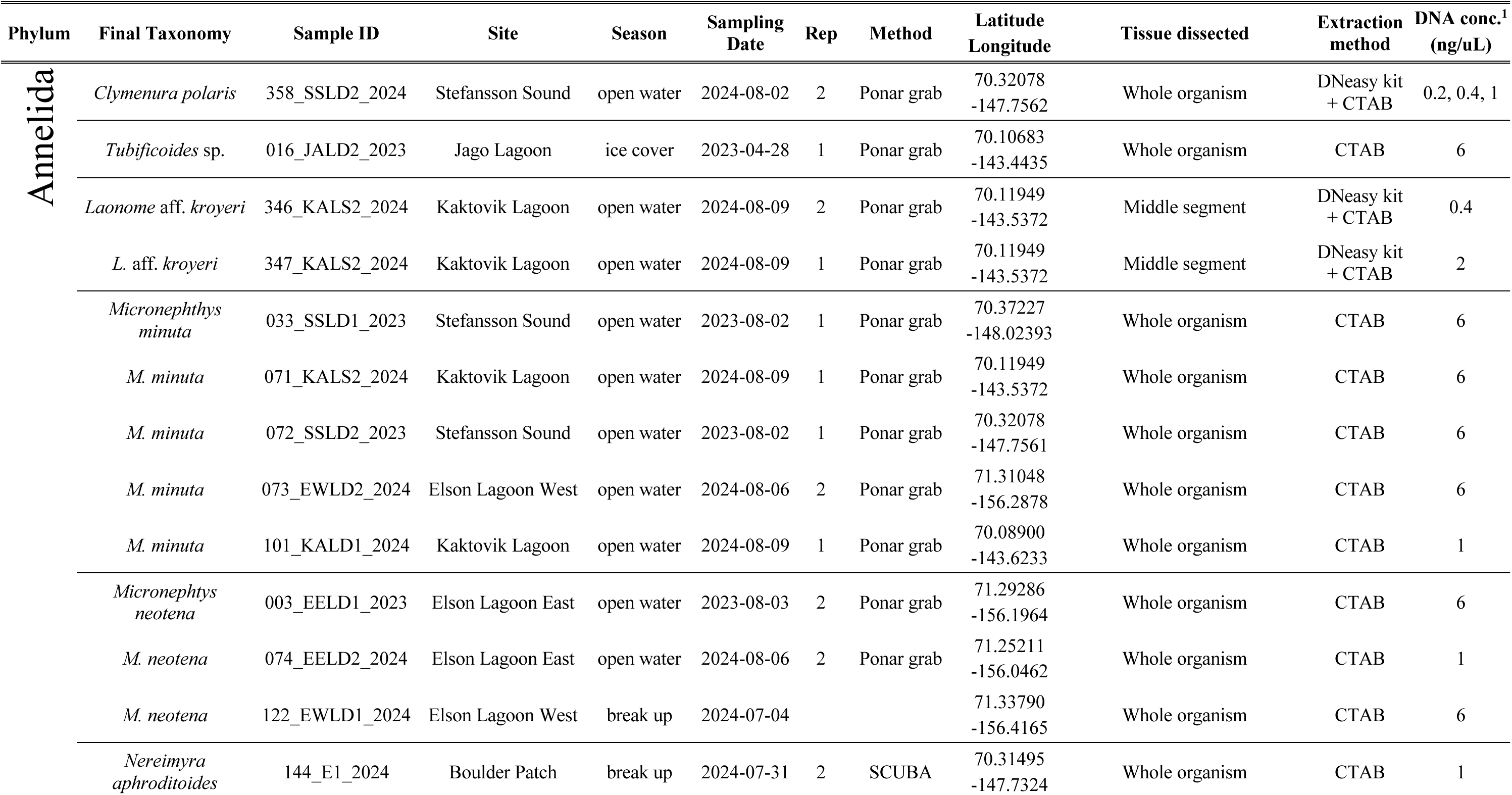
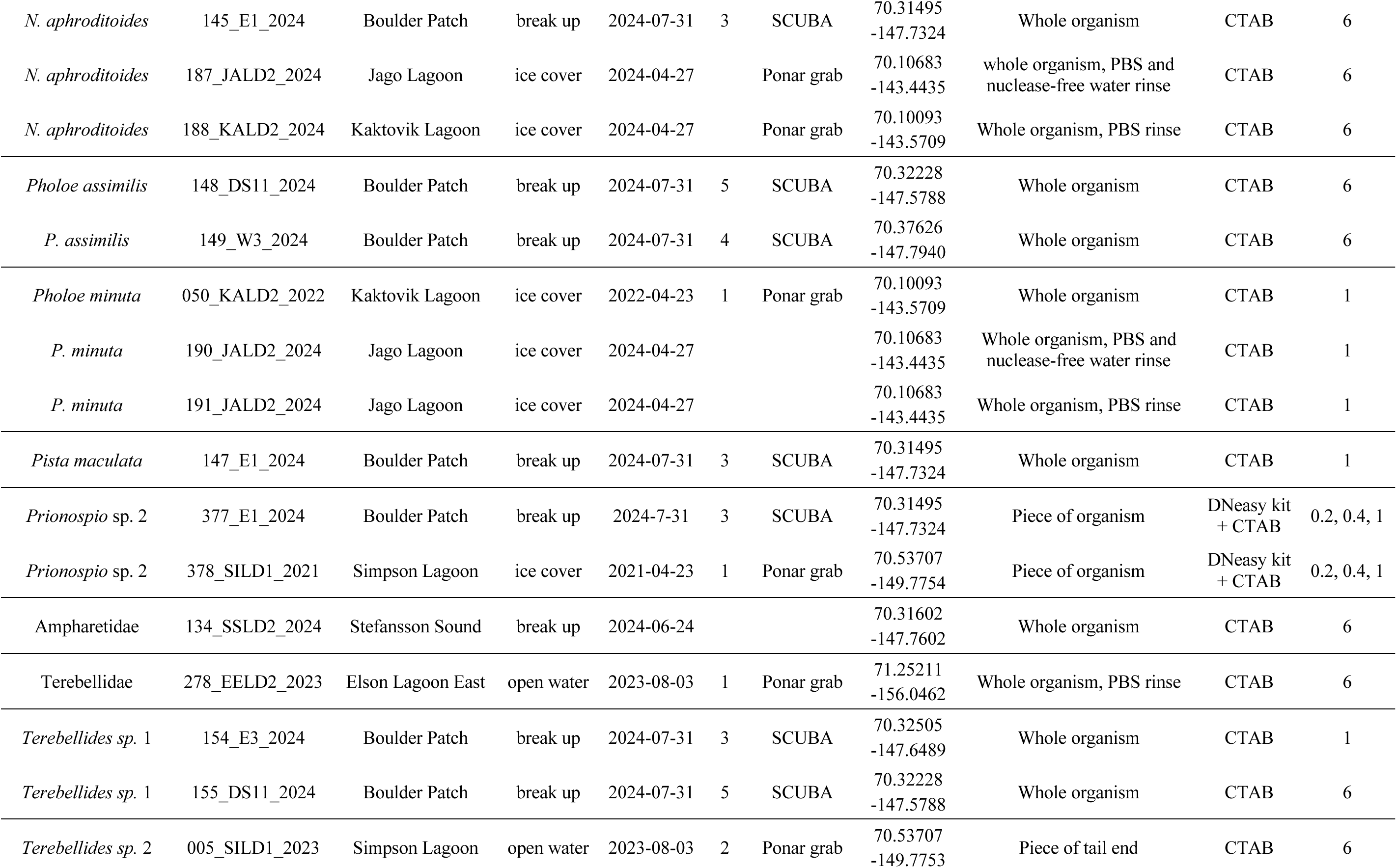
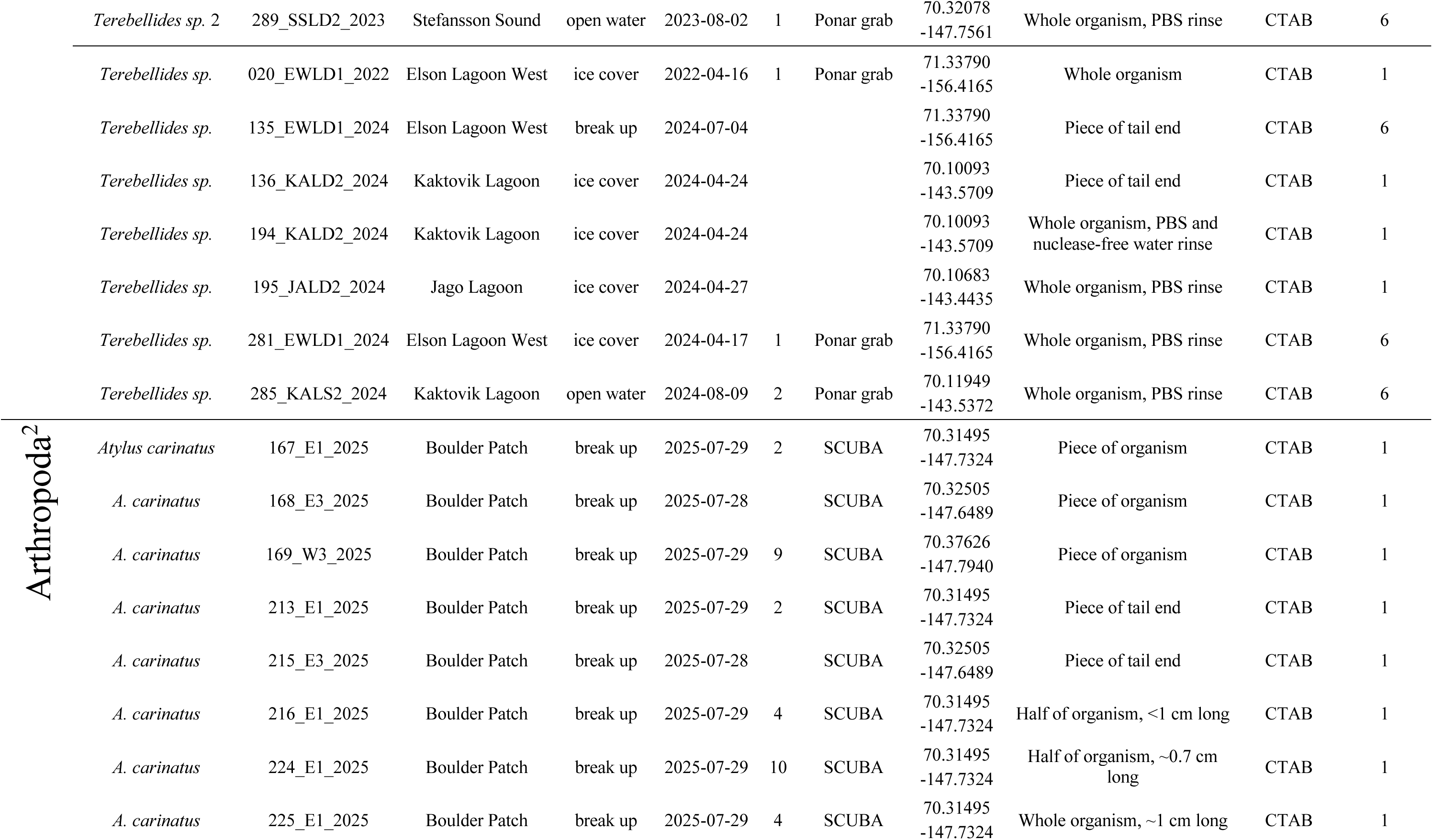
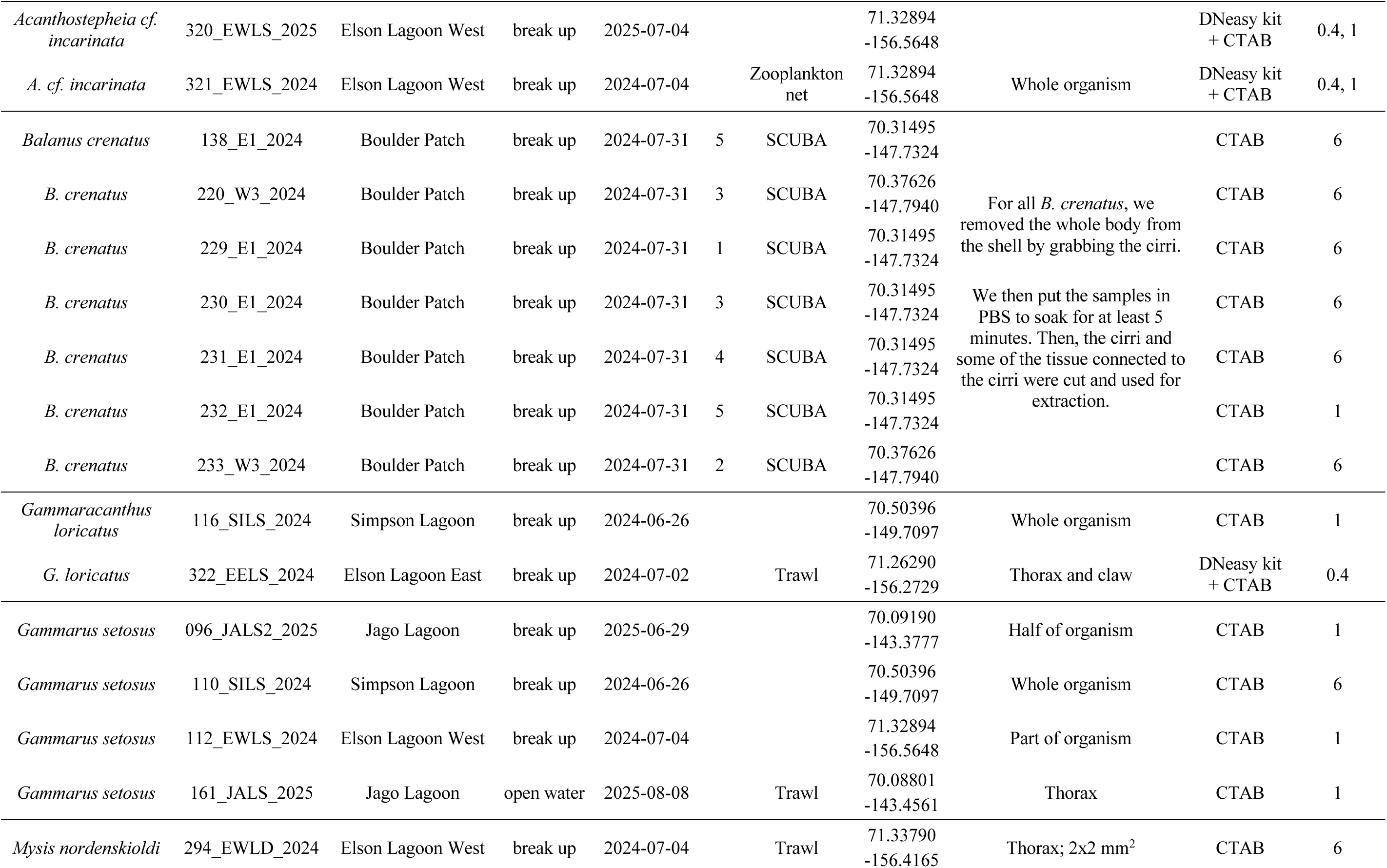
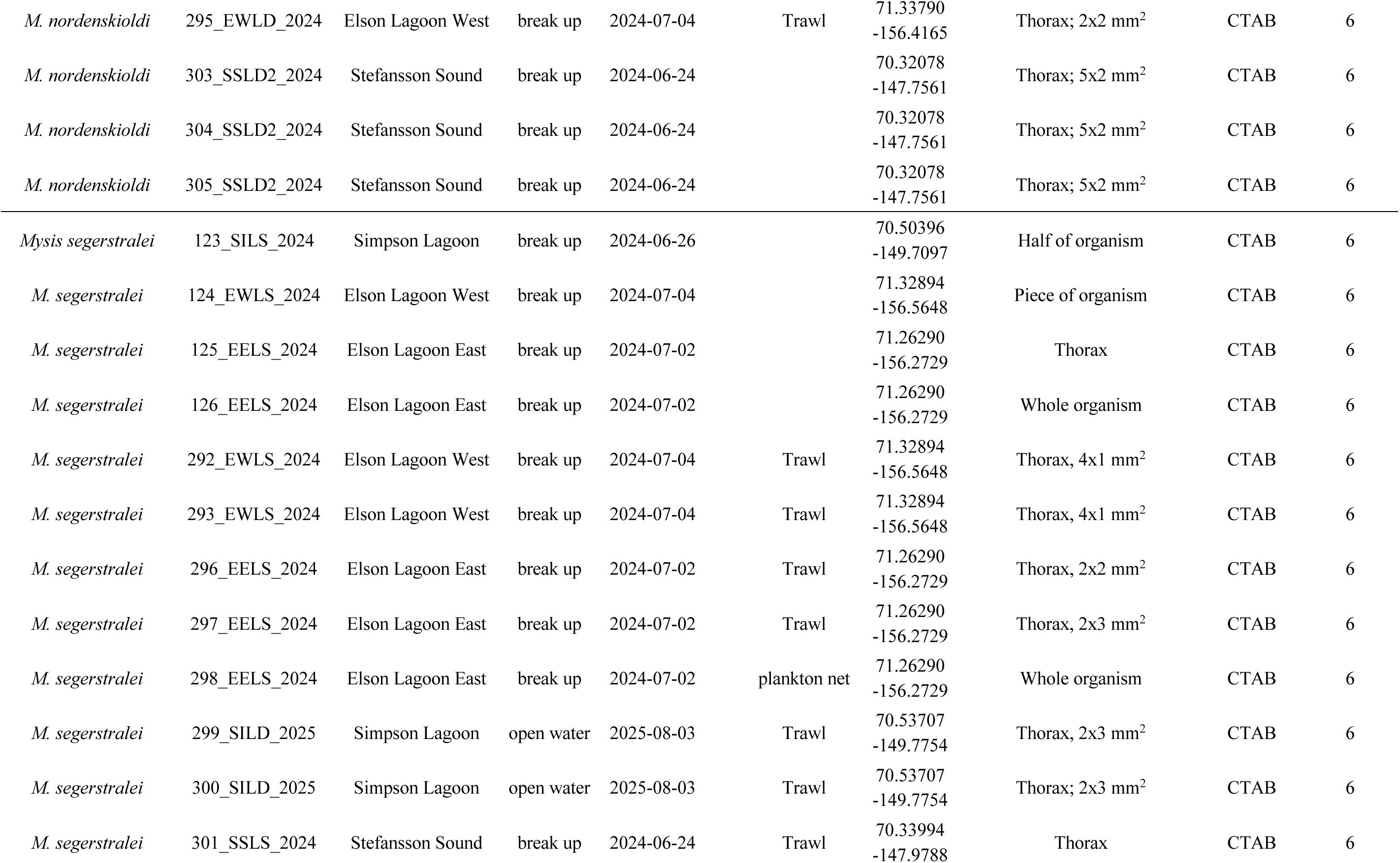
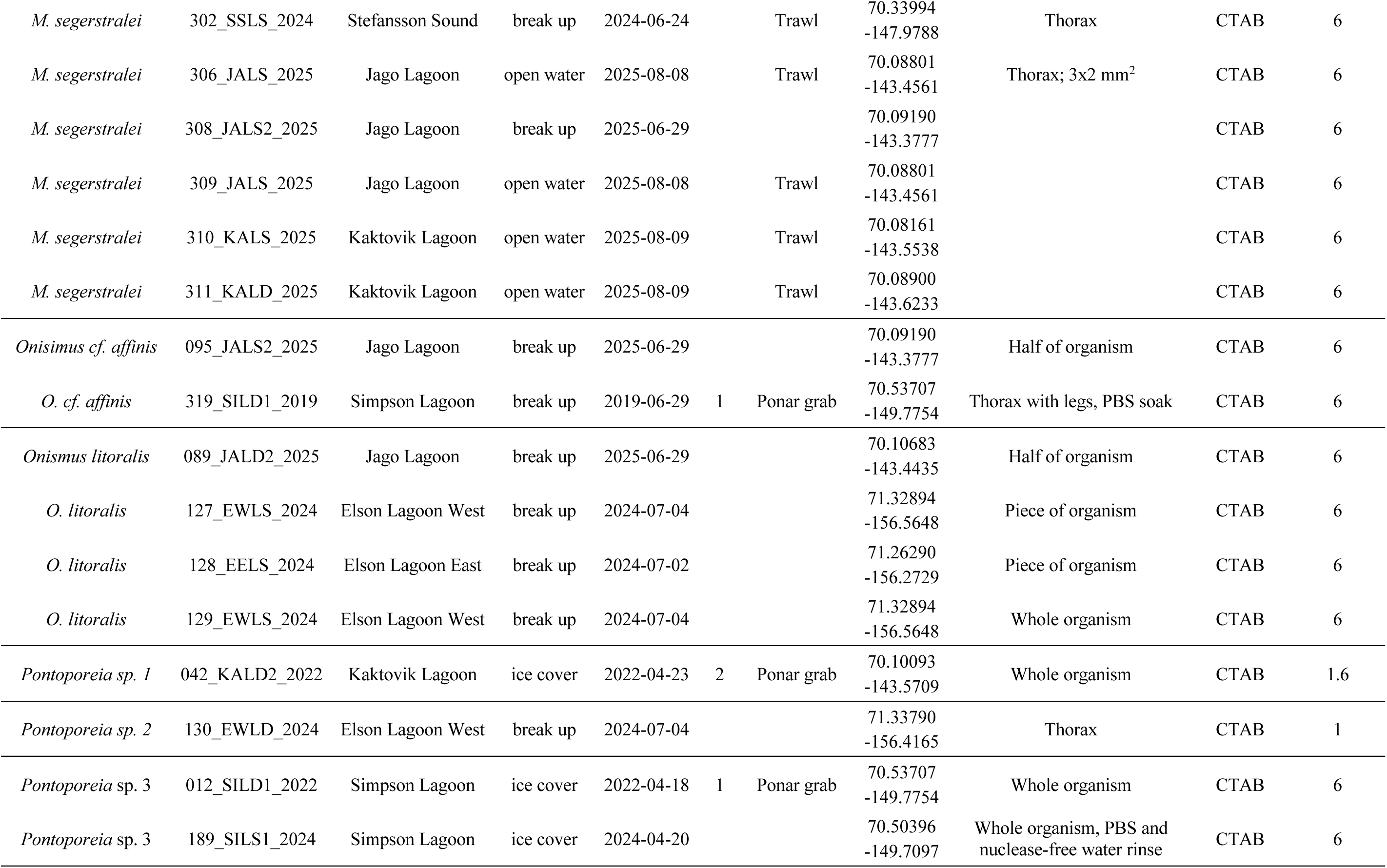
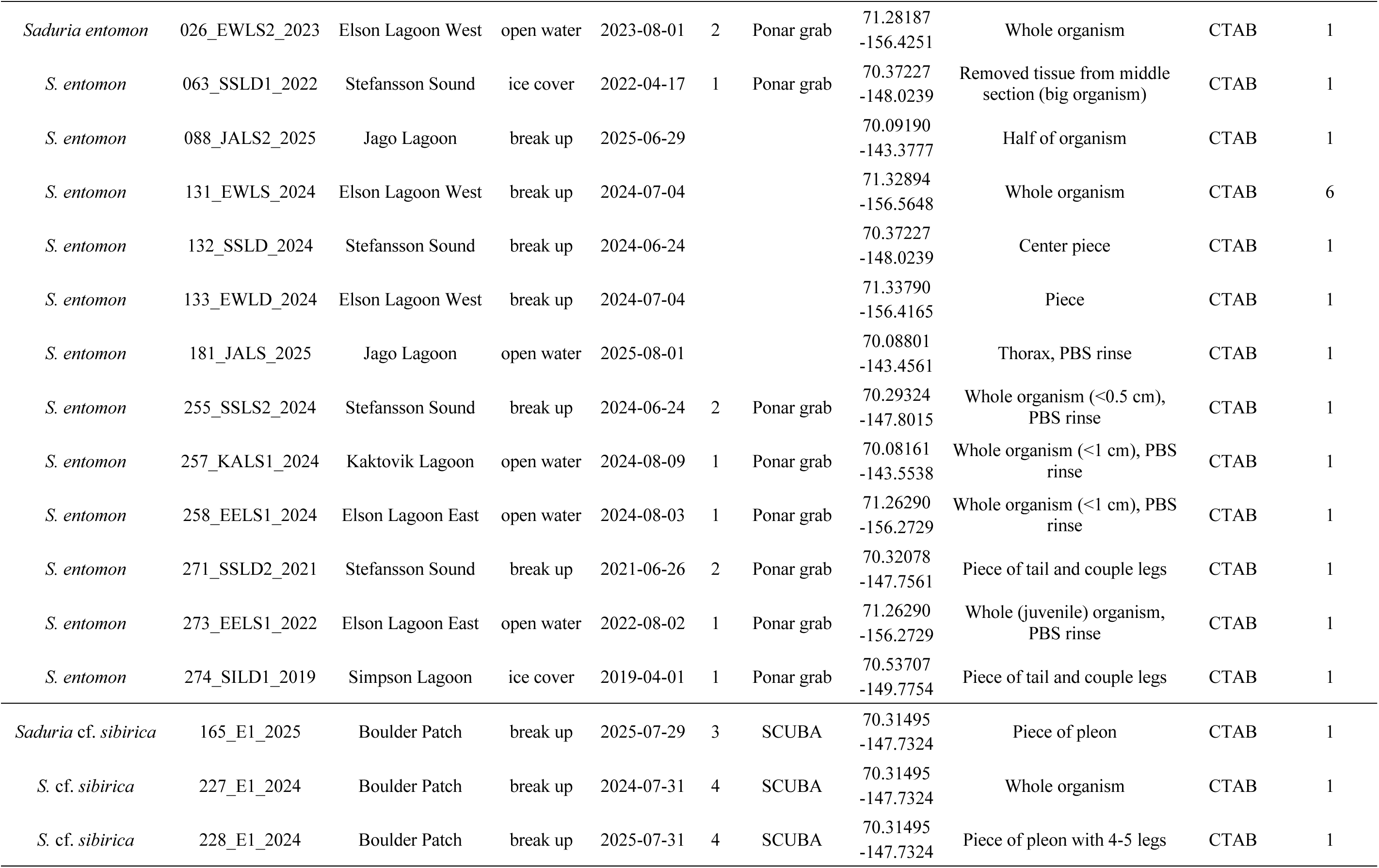
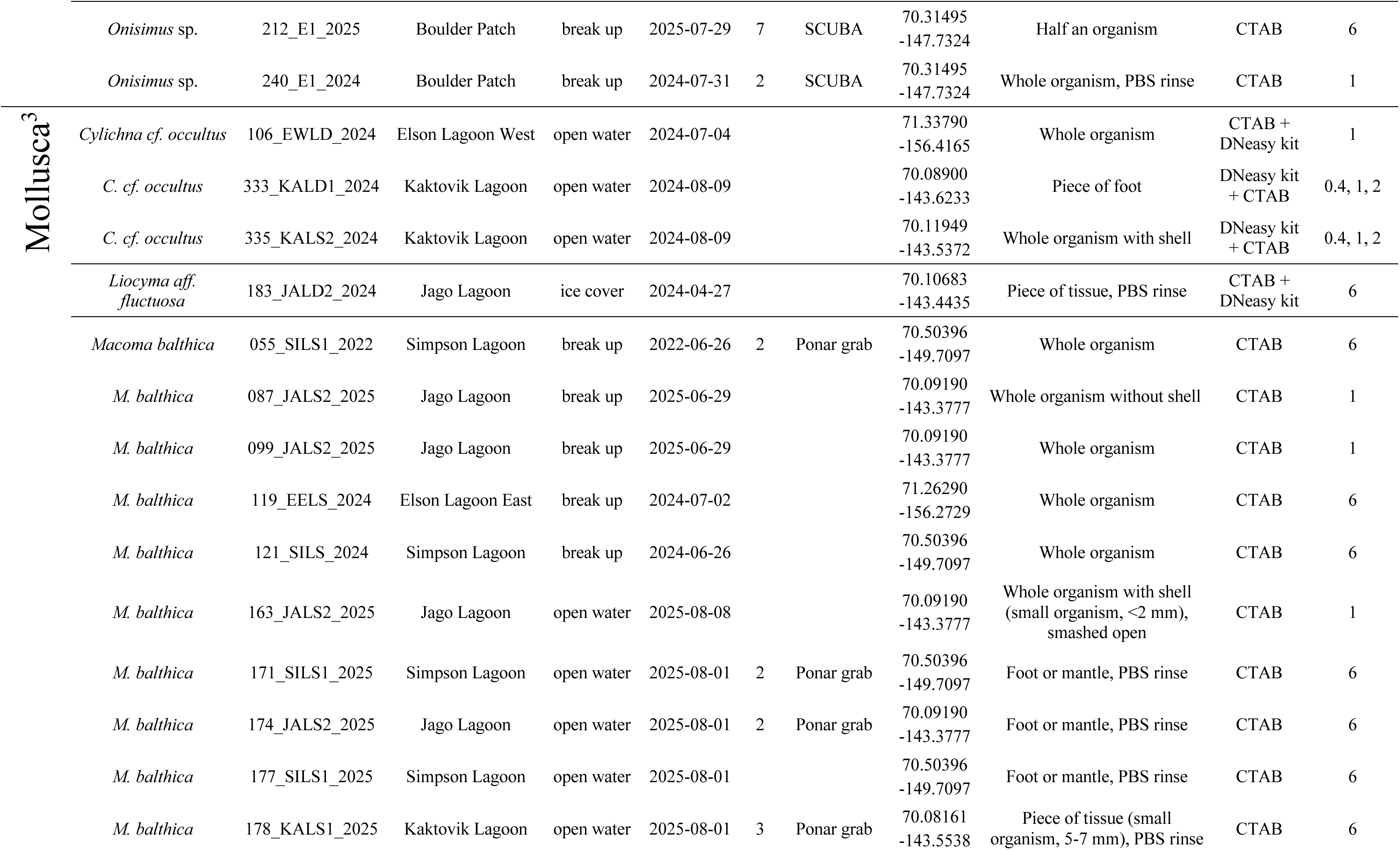
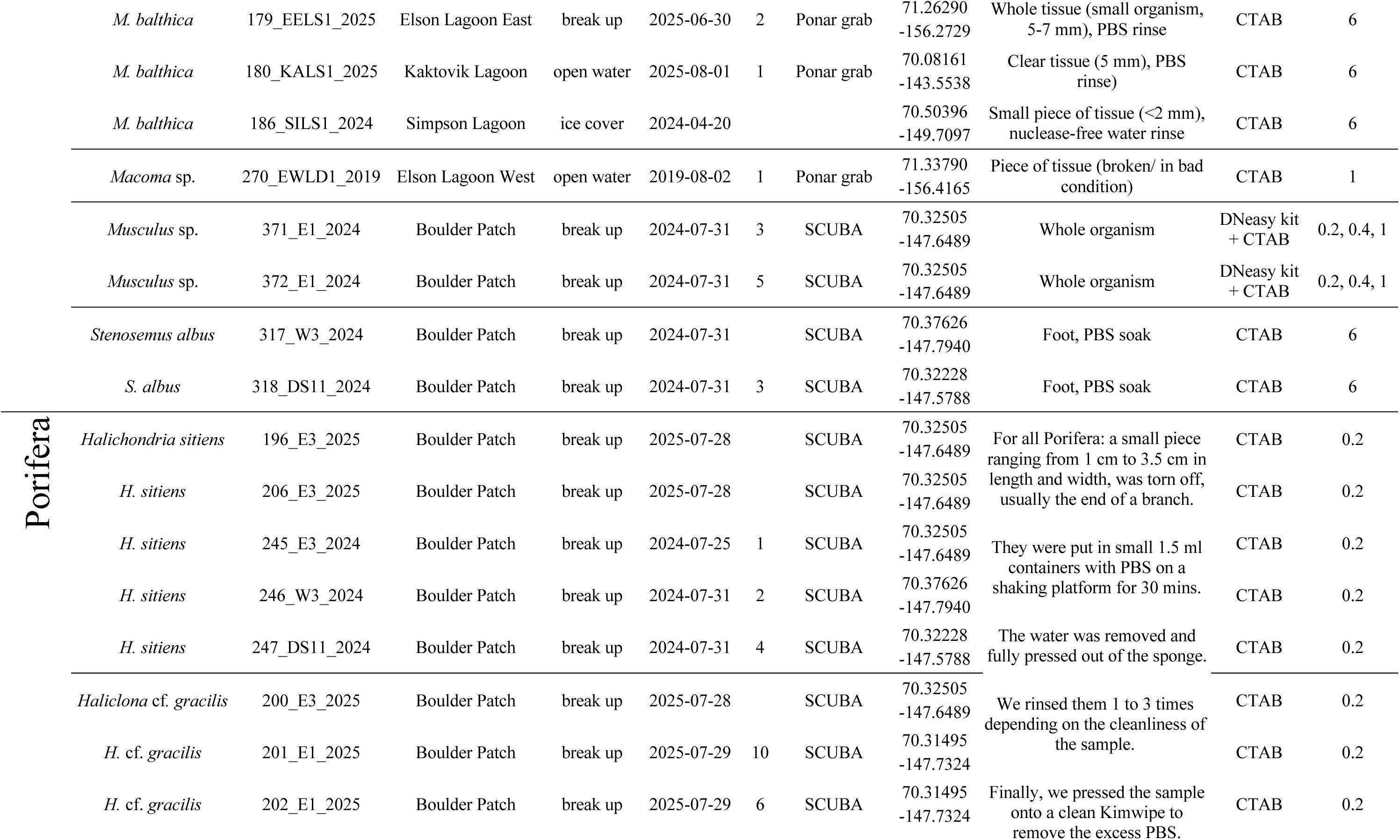
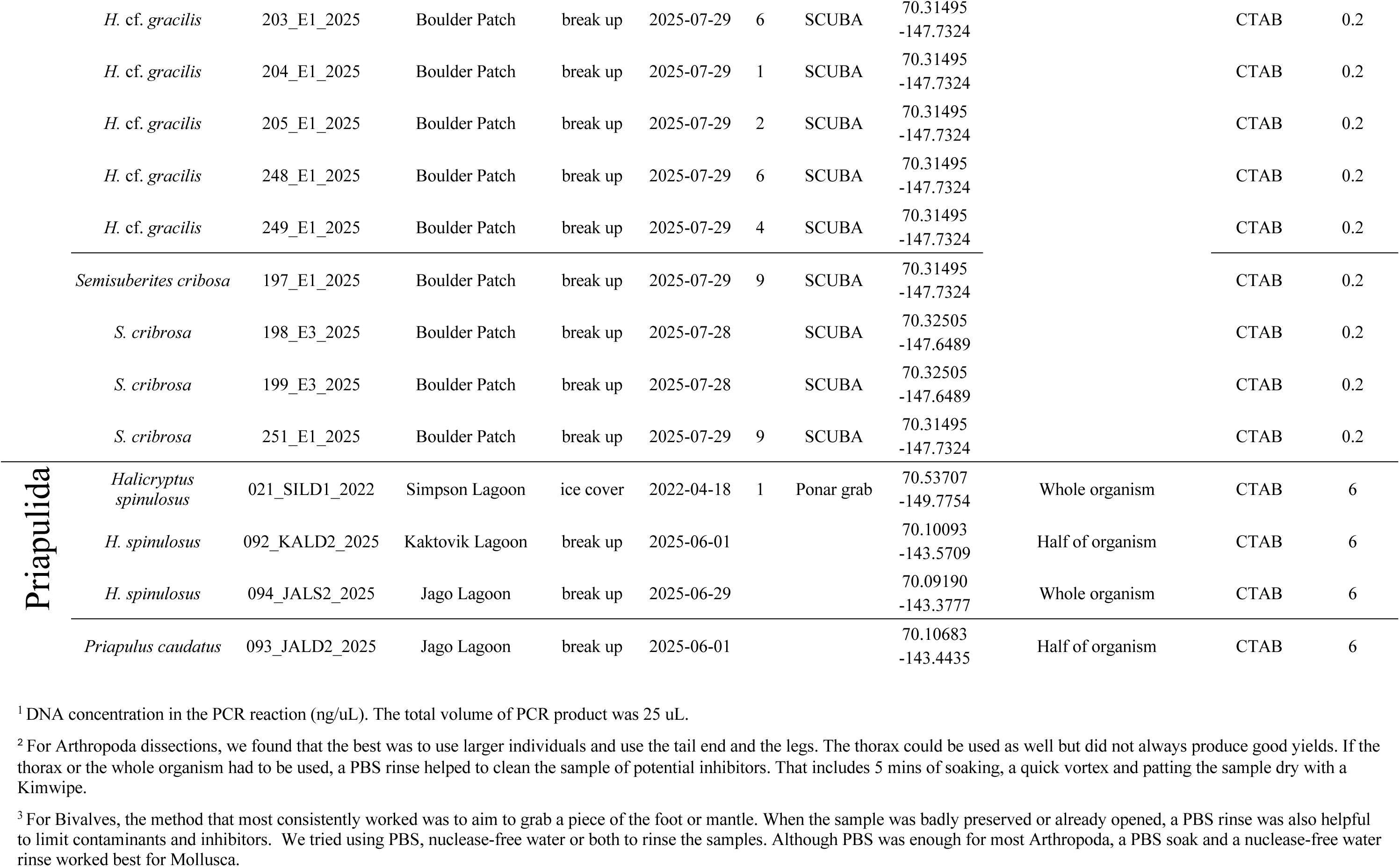
Final taxonomy based on morphological and molecular identification for each individua site, season, date (YYYY-MM-DD), sample replicate (Rep) if applicable, sampling method if known, and coordi Molecular method details: tissue dissected from each individual if known, extraction method used, DNA concent used for amplification (if multiple concentrations are listed they all worked and were combined for sequencing).

